# A single-nucleus atlas of the adult laying hen liver reveals metabolic specialization and improved cellular resolution through enhanced genome annotation

**DOI:** 10.64898/2026.07.27.740890

**Authors:** Laetitia Lagoutte, Coralie Allain, Bénédicte Lebez, Guillaume Cossard, Frédéric Lecerf, Yuna Blum, Sandrine Lagarrigue, Fabien Degalez

## Abstract

The liver of laying hens plays a central role in metabolism and reproduction, supporting the synthesis of egg yolk precursors under strong hormonal regulation. Despite its physiological importance, a high-resolution cellular reference of the adult chicken liver is still lacking. Here, we generated a single-nucleus RNA sequencing atlas of the adult laying hen liver from eight individuals, providing a comprehensive view of its cellular composition and transcriptional landscape.

Using this framework, we identified major hepatic cell populations, including hepatocytes, endothelial cells, cholangiocytes, hepatic stellate cells, and diverse immune cell types, revealing a broadly conserved vertebrate liver architecture. However, hepatocyte zonation, a key feature of mammalian liver organization, was not observed, consistent with the absence of hepatocyte zonation reported in birds.

Importantly, we demonstrate that the use of an enriched genome annotation, incorporating additional protein-coding and long non-coding RNA models, substantially improves transcript detection and enhances cell-type resolution in single-nucleus datasets. This improved resolution allows more accurate marker-based assignment of hepatocyte subpopulations and refines the interpretation of hepatic cellular heterogeneity.

Within hepatocytes, we uncovered transcriptionally distinct subpopulations associated with lipid metabolism and reproductive function, including estrogen-responsive programs involving cytochrome P450 genes such as CYP2C23A and CYP2C23B. In parallel, we characterized a complex immune compartment composed of resident macrophages and adaptive immune cells, highlighting the dual metabolic and immunological roles of the avian liver.

Overall, this atlas provides a high-resolution reference for avian liver biology and demonstrates that improved genome annotation enhances the resolution and interpretation of cellular heterogeneity in single-cell transcriptomic studies.

## INTRODUCTION

The chicken is a major livestock species supplying protein-rich food worldwide, with global poultry meat production exceeding 138 million tonnes and egg production reaching around 91 million tonnes in 2024 (1,2). Beyond its economic importance, the chicken is also a widely used model organism in both fundamental and applied research, contributing to our understanding of development (3), immunology (4), host–pathogen interactions (5), and evolutionary biology (6). Among chicken organs, the liver is of particular interest because of its central roles in metabolism, detoxification, and immune regulation. In laying hens, hepatic function is further specialized to support reproduction through the synthesis of yolk precursors, such as very-low-density lipoproteins (VLDL) and vitellogenins, which are essential for egg formation (7,8). These combined metabolic and reproductive demands make the adult chicken liver a uniquely informative system for studying cellular heterogeneity and physiological regulation.

Over the past decade, bulk RNA sequencing has been extensively used to investigate gene expression in the chicken liver under diverse experimental conditions providing insights into metabolic regulation (9), immune responses (10), and egg production (11). Consequently, liver samples represent a consequent proportion of chicken RNA-seq datasets available in public repositories. Large-scale initiatives such as FAANG (12,13) and FarmGTEx (14) have generated extensive transcriptomic resources for livestock species and the recent ChickenGTEx project (15,16) has further expanded this landscape by integrating bulk RNA-seq with genetic and epigenomic data across multiple tissues including liver. While these efforts have greatly advanced our understanding of hepatic gene regulation, bulk transcriptomic approaches capture only average expression profiles and cannot resolve the cellular composition or cell-type–specific transcriptional programs underlying liver function.

The liver is composed of parenchymal cells, including hepatocytes and cholangiocytes, as well as multiple non-parenchymal populations such as endothelial cells, immune cells, and mesenchymal cells. Historically, these populations have been characterized through histological analyses, flow cytometry, and selected molecular marker based approaches that provided important foundational knowledge but lacked cellular resolution at the transcriptomic level. Single-cell and single-nucleus RNA sequencing (scRNAseq / snRNAseq) now enables systematic dissection of liver cellular diversity and functional states (17). In mammals, such approaches have fundamentally reshaped liver biology, revealing hepatocyte zonation (18,19), endothelial specialization, and diverse immune-cell population (20). Comparative studies suggest that functional hepatocyte zonation has been best characterized in mammals (21) .

In avian species, however, high-resolution cellular maps of the liver remain limited. A recent cross-species single cell study by Yuan et al., 2025 (22) demonstrated that birds, including chickens, exhibit distinct transcriptional programs and regulatory architectures, with lipid metabolism genes evolving rapidly, highlighting the need for species-specific high-resolution analyses.

To date, despite studies such as those by Yuan et al., 2025 (22), which adopt a multi-species approach, and chickenGTEx (15,16), which integrates single-cell analyses into a broader research framework, single-cell studies in avian species remain limited. In hens more precisely, snRNAseq has mainly been applied to embryonic and newly hatched chicken livers (23), where major cell populations and developmental changes in lipid metabolism have been described. While informative for liver development, this study does not address the adult stage, where hepatic function is profoundly shaped by the metabolic demands associated with egg production. In contrast, comparable cellular atlases are now emerging in other livestock species, such as cattle (24) and pig (25) emphasizing the current lack of a comprehensive reference for the adult chicken liver.

Accurate genome annotation also represents a critical challenge for single-cell transcriptomics in non-model organisms. Incomplete or fragmented gene models can directly affect transcript detection, cell-type identification, and downstream biological interpretation. Recent improvements in the chicken genome assembly, notably the GRCg7b release, have incorporated microchromosomes and refined previously characterized chromosomes (26). In parallel, independent annotation efforts have improved protein-coding gene models and identified thousands of long non-coding RNA genes (27,28), providing an opportunity to reassess in a second time, hepatic cellular diversity at higher resolution and to evaluate how annotation strategies influence single-cell analyses.

To address these gaps, we performed single-nucleus RNA sequencing on eight frozen liver samples from adult laying hens to generate a comprehensive atlas of hepatic cell populations. In addition to defining the major hepatic cell populations and transcriptional programs, we assessed how genome annotation strategy influences cell-type resolution and biological interpretation in avian single-cell datasets. Together, this work provides a high-resolution cellular reference for the adult female chicken liver and establishes a framework for investigating hepatic metabolism, immunity, and reproductive specialization in birds.

## MATERIAL AND METHODS

### Animals and sample collection

The study involved Rhode Island Red pure-bred hens who were raised according to commercial standards with *ad libitum* commercial laying diet, and kept under a 16-hour light/8-hour dark cycle at an ambient temperature of 20°C. Layers were slaughtered at 90 weeks of age at the fed status by neck cut and bleeding, immediately after head electrical stunning. Right after slaughter the extremity of the left liver lobe was sampled, snapped frozen in liquid nitrogen and stored at -80°C until nuclei isolation. The experimental protocol complied with animal welfare standards and received approval from the ethics committee for animal experimentation (APAFIS #34628-2022011214434077 v3) in agreement with the French and European legislations.

### Nuclei suspension preparation

Nuclei were isolated using the Tissue Nuclei Extraction for ATAC-seq kit (C01080003, Diagenode). Briefly, 50 mg of frozen liver were transferred to a Dounce homogenizer containing 800 µl of tissue lysis buffer supplemented with 4 µl Rnase inhibitor (Applied biosystems *#N8080119*). Tissues were disrupted with the loose pestle (A) for 20 strokes followed by the tight pestle (B) until a homogenous suspension was obtained. The suspension was filtered twice through a 30µm cell strainer (*MACS smart strainer (Miltenyi Biotec #130-098-458)*. Homogenates were then loaded onto an iodixanol gradient and centrifuged at 3,500 × g for 20 minutes at 4°C using a swing-out rotor. A thin whitish band corresponding to nuclei was collected between layers 2 and 3 and observed in confocal microscopy with Trypan Blue to assess nuclear integrity. Nuclei were counting using the LUNA-FL automated counter and the AO/PI kit (Logos biosystems *#*F23001).

### Library preparation and sequencing

Nuclei fixation and library preparation were performed following the manufacturer’s instructions (Parse Biosciences, Evercode Nuclei Fixation v3 and Evercode WT v3). cDNA library quantity and quality were assessed using the Qubit dsDNA HS Assay kit (Invitrogen, *#*Q32851) and the High Sensitivity DNA kit (Agilent *#*5067-4626). Libraries were sequenced on an Illumina Novaseq X plus platform using 100 bp paired-end sequencing, targeting approximately 50,000 read pairs per nucleus.

### Data processing and quality control

Raw FASTQ files were processed using the TrailMaker™ pipeline (Parse Biosciences, v1.5.0; https://app.trailmaker.parsebiosciences.com/) with default parameters and aligned to the chicken reference genome GRCg7b (GCF_016699485.2). Two annotations were used : *i)* the reference annotation from Ensembl v114 initially including 17,007 protein-coding genes (PCGs) and 11,944 long non-coding RNA genes (lncRNAs) (https://ftp.ensembl.org/pub/release-114/gtf/gallus_gallus/Gallus_gallus.bGalGal1.mat.broiler. <u>GRCg7b.114.chr.gtf.gz</u>) and *ii)* the gene-enriched annotation from Degalez et al., 2024 (27) including 24,102 PCGs and 44,428 lncRNAs. This atlas is also available through the GEGA interface (https://gega.sigenae.org/download) (28).

For both annotations, only PCGs and lncRNAs were retained and genes located on the mitochondrial chromosome were also excluded. This results in final annotations comprising, respectively i) 16,994 PCGs and 11,944 lncRNAs for Ensembl annotation and ii) 23,900 PCGs and 44,428 lncRNAs for the gene-enriched annotation respectively.

For both annotations, the resulting unfiltered gene–cell count matrix and associated metadata were downloaded for downstream analyses and imported into R via the *Seurat* package (v5.3.0 ; (29,30)) using the *ReadParseBio* function. These files were initially used to generate two Seurat objects : *i*) An unfiltered object (min.features = 0, min.cells = 0) to assess raw data distributions and ; *ii*) A pre-filtered object (min.features = 200, min.cells = 3) to remove low-quality nuclei and lowly detected genes.

Subsequently, quality control metrics, including the number of detected genes per nucleus (*nFeature_RNA*) and the total number of detected transcripts (*nCount_RNA*), were examined for each sample independently using violin plots and feature scatter plots (Supplementary Figure 1A). To account for sample-specific variability in sequencing depth and transcript detection, filtering was performed independently for each sample, using a filtering strategy adapted from the Trailmaker pipeline. For each sample, empirical upper thresholds were defined as the 95th percentile of the distributions of *nFeature_RNA* and *nCount_RNA* (Supplementary Table 1). Nuclei exceeding either threshold were excluded. Filtered datasets from all samples were then merged into a single Seurat object (*JoinLayers*).

### Data integration and normalization

Data were normalized using the *NormalizeData* function with the *LogNormalize* method and a scale factor of 10,000. Highly variable genes (HVGs) were identified using the variance stabilizing transformation (VST) method implemented in *FindVariableFeatures*, selecting the top 2,000 genes (Supplementary Figure 1B). HVGs were so computed on the unified dataset to capture global biological variability across all nuclei. Data were then centered and scaled with the *ScaleData* function.

### Dimensionality reduction

Principal component analysis (PCA) was performed on the scaled expression matrix using the selected highly variable genes, reducing dimensionality to 30 components (Supplementary Figure 1C).

### Batch effect correction

To correct for inter-sample technical variability, batch effects were addressed using the Harmony algorithm (Harmony v1.2.3; (31,32)) and with the sample identity as grouping variable. Harmony was applied to the PCA embedding, generating a corrected low-dimensional representation of the data. Convergence diagnostics were monitored to ensure stable optimization.

### Non-linear dimensionality reduction and visualization

Uniform Manifold Approximation and Projection (UMAP) was computed using the Harmony-corrected embedding with the cosine distance as similarity metrics and a minimum distance parameter of 0.3. UMAP embeddings were visualized to assess sample mixing and overall data structure.

### Cluster identification

Cell–cell nearest-neighbor graphs were constructed using the Harmony-corrected dimensions via the *FindNeighborsfunction*. Clustering was performed using the Leiden algorithm (algorithm = 4) implemented in *FindClusters*.

Multiple resolution parameters (0.1, 0.3, 0.5, 0.8, and 1.0) were evaluated to assess clustering granularity. Cluster stability and hierarchical relationships across resolutions were examined using the *clustree* package (v0.5.1; (33,34)). A resolution of 0.1 was selected for downstream analyses of major cell populations.

### Identification of cluster-specific marker genes and cluster assignation

Cluster-specific marker genes, hereafter referred to as differentially expressed genes (DEGs), were identified using the Wilcoxon rank-sum test as implemented in the presto package (v1.0.0; (35,36)) through the wilcoxauc function. For each cluster, gene expression levels were compared against all remaining cells. Genes were subsequently ranked according to their area under the curve (AUC), which reflects their discriminatory power by integrating both the magnitude of differential expression (fold change, FC) and the proportion of cells expressing the gene within the cluster relative (pct-in) to all other cells (pct-out). The top-ranked marker genes (typically the top 10 per cluster) were then compared with previously reported cell type–specific markers from the literature (Supplementary Table 2) and publicly available databases to support cluster annotation.

### Subclustering analysis

To further refine cellular heterogeneity within major liver populations, subclustering analyses were independently performed on selected macro-clusters identified during the initial global analysis. Cells belonging to the corresponding clusters were extracted from the integrated Seurat object using cluster identities obtained at the macro-clustering resolution. This was performed for hepatocyte, endothelial and immune cells.

The subclustering workflow followed the same analytical strategy as the global analysis. Briefly, subsetted datasets were normalized using the *LogNormalize* method with a scaling factor of 10,000. Highly variable genes were identified using the variance stabilizing transformation (*vst*) method implemented in Seurat, retaining the 2,000 most variable features. Data were subsequently scaled prior to dimensionality reduction by Principal Component Analysis (PCA), computed on the first 30 principal components.

To minimize inter-sample technical variability while preserving biological structure, Harmony integration was applied using the sample identifier as batch covariate. Uniform Manifold Approximation and Projection (UMAP) was then performed on the Harmony embeddings using cosine distance and a minimum distance parameter of 0.3 for visualization of cellular relationships.

Cellular neighborhoods were identified using the shared nearest neighbor (SNN) graph computed from the Harmony reduction, and clustering was performed using the Leiden algorithm (algorithm 4 in Seurat). Multiple clustering resolutions (0.1, 0.3, 0.5, 0.8 and 1.0) were explored to evaluate the stability and granularity of subpopulation structures. Resulting subclusters were subsequently annotated based on the expression of known marker genes and differential expression analyses.

### Cell cycle index estimation

Cell cycle phase assignment was performed using the *CellCycleScoring* function implemented in Seurat. Cells were classified into G1, S, and G2/M phases based on canonical marker genes provided in the Seurat package. Given species-specific differences and incomplete annotation in chicken, the gene lists were manually curated to ensure compatibility with the *Gallus gallus* genome. For the G2/M gene set (54 genes in the original list), 49 genes were directly matched to chicken orthologs. The five missing genes were replaced as follows: *UBE2C* by *UBE2U* (1:1 ortholog), *CCNB2* by *CCNB1* (1:1 ortholog and synonym), *CDC25C* by multiple predicted orthologs (ENSGALG00010029451, ENSGALG00010029468, ENSGALG00010029469, ENSGALG00010029471, ENSGALG00010029472, ENSGALG00010029474, ENSGALG00010029475, ENSGALG00010029476", ENSGALG00010029942 ; 1:many orthology relationships), *CDCA8* by *CDC25A* (based on pairwise genomic alignment), and *AURKA* by *AURKB* (1:1 ortholog). For the S phase gene set (43 genes), only one gene (*MLF1IP*) was absent and was replaced by *CENPU*, based on gene name synonymy.

### Function enrichment analysis

Functional enrichment analysis was performed to identify overrepresented Gene Ontology (GO) biological processes among cluster-specific marker genes (37,38). For each cluster, only genes with an adjusted p-value < 0.05 were retained, and the top 50 genes (ranked by AUC) were selected for enrichment analysis. Enrichment was conducted using the *GO_Biological_Process_2025* database via the *enrichR* package (v3.4; (39,40)), and only pathways with an adjusted p-value < 0.05 were considered significant. All enrichment results are provided in Supplementary Table 3. An additional summary table (Supplementary Table 4) reports, for each cluster, the number of significant marker genes, the number of genes used for enrichment analysis (capped at 50), and associated statistics for these genes regarding the AUC, logFC and number of unnamed genes.

### Cluster stability across annotations

To assess cluster stability between the two annotations used, both at the global level and within lineage-specific subclusters, we computed the Adjusted Rand Index (ARI) using the *adjustedRandIndex* function from the *mclust* (v6.1.2; (41,42)) package. This metric quantifies the agreement between two clustering solutions while correcting for chance, thereby providing a robust measure of clustering consistency across annotation frameworks. An ARI value of 1 indicates perfect concordance between clusterings, whereas a value of 0 corresponds to random agreement

## RESULTS

### ATLAS OF SINGLE NUCLEI IN THE LIVER OF LAYING HENS

#### General statistics and data processing

Single-nucleus RNA sequencing (snRNA-seq) was performed on liver samples from eight adult laying hens (Figure 1A), yielding a total of 2,112,583 nuclei prior to filtering. After quality-control filtering based on the distributions of the number of genes and transcript detected (Supplementary Table 1), 117,263 nuclei were retained for downstream analyses. The median number of nuclei recovered per sample was 9,401, with a broadly balanced representation across six of the eight samples. In contrast, samples C122 and C140 contributed larger numbers of nuclei, with 38,249 and 24,960 nuclei, respectively. Sequencing depth was overall homogeneous across nuclei and across samples, with a median of approximately 6,968 reads per nucleus. The median number of genes detected per nucleus was 862 among the 28,938 genes in Ensembl annotation, showing limited inter-sample variation. Likewise, the mean number of transcripts per nucleus ranged from 1,104 in sample C62 to 2,142 in sample C140, with an overall median of 1,561 across samples, indicating robust and consistent transcript capture throughout the dataset.

**Figure 1.**
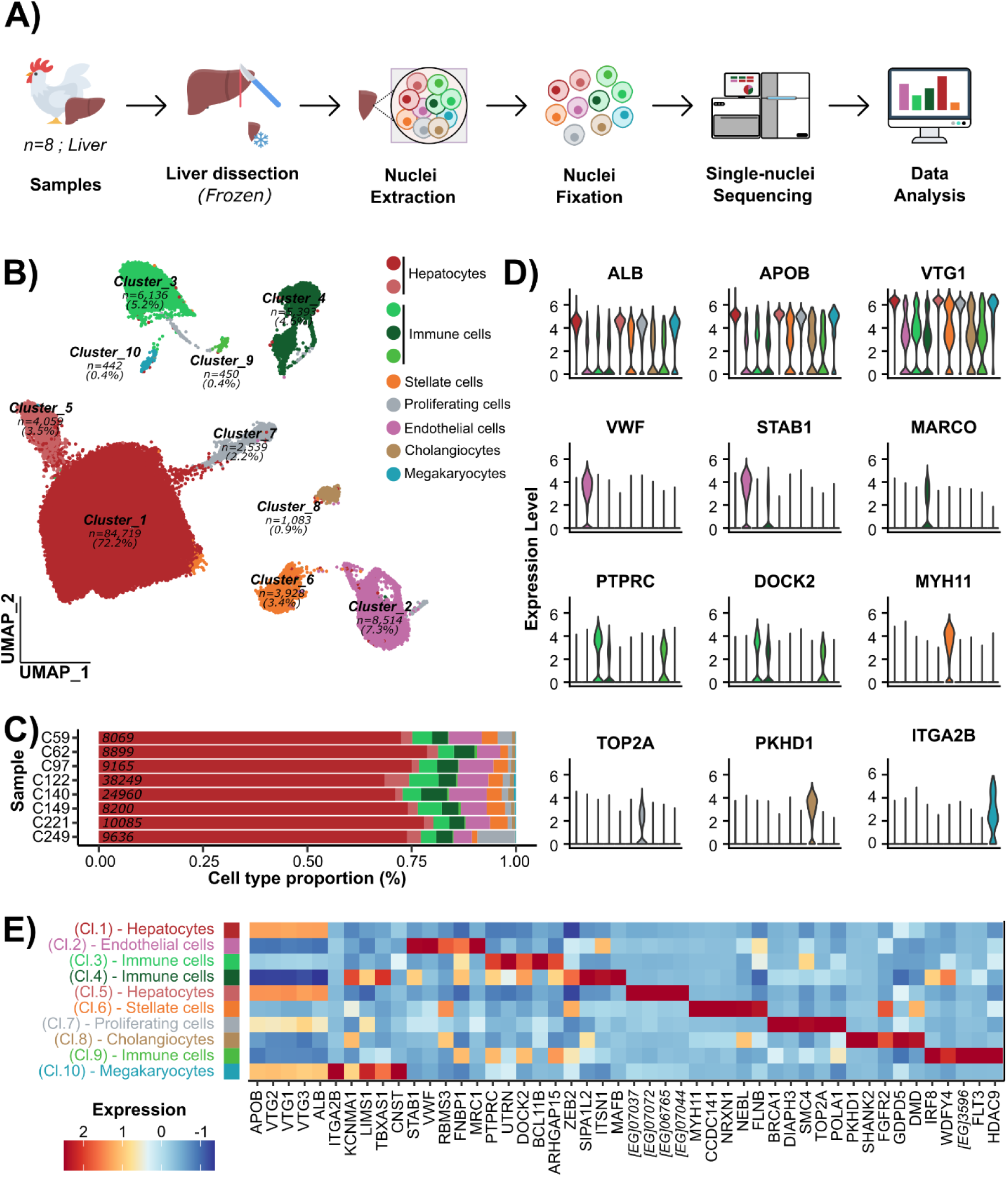
Single-nucleus RNA-seq profiling reveals the cellular composition of major cell types in the adult chicken liver. **(A)** Overview of the single-nucleus RNA-sequencing analysis from sampling to data analysis. **(B)** UMAP of integrated single-nucleus RNA-seq data highlighting ten transcriptionally distinct clusters (indicating in italics) identified by unsupervised clustering (Leiden algorithm, resolution = 0.1). Cluster biological identities were assigned based on marker genes identified in panels (D) and (E). (n=) indicates the number of cells associated with the cluster and the corresponding percentage **(C)** Proportion of each major cell type across the eight individual samples; numbers in italics indicate the total number of nuclei per sample. **(D)** Distribution of gene expression levels for selected canonical marker genes across clusters. **(E)** Top five differentially expressed genes per cluster (one-versus-all), supporting the identification of major hepatic cell populations. Cl.: Cluster ; [EG]: ENSGALG000100.

### Clustering and integration of single-nucleus transcriptomes

Unsupervised clustering of the integrated single-nucleus RNA-seq dataset was performed across the eight samples using Harmony-corrected principal components and visualized by UMAP (Supplementary Figure 1E-F). Nuclei from all samples were well mixed in the low-dimensional embedding, with no detectable sample-specific segregation, despite two samples (C122 and C140) contributing approximately four- and two-fold more nuclei, respectively, than the others.

To assess clustering stability and resolution-dependent granularity, unsupervised clustering was performed across a range of resolutions (0.1, 0.3, 0.5, 0.8, and 1.0) using the Leiden algorithm (Supplementary Figure 2A). Increasing the resolution progressively subdivided the dataset into 10, 13, 17, 21, and 23 clusters, respectively. Examination of cluster structure and continuity across resolutions indicated that a resolution of 0.1 provided a global and interpretable representation of the expected major cell populations. This resolution was therefore selected for subsequent global analyses (Figure 1B). At this resolution, the proportion of nuclei assigned to individual clusters ranged from 0.4% (Cluster 10) to 72% (Cluster 1), reflecting the expected predominance of potential hepatocytes alongside rarer non-parenchymal populations (Figure 1C).

### Annotation of Major Hepatic Cell Populations

Putative cell identities were assigned to clusters based on the expression of canonical lineage-specific markers across clusters (Figure 1D; Supplementary Table 2), together with differential gene expression analyses highlighting distinct transcriptional programs (Figure 1E).

Based on canonical markers, this analysis identified seven major lineages expected in the liver: hepatocytes (clusters 1 and 5; **ALB** marker), endothelial cells (cluster 2; **VWF**), cholangiocytes (cluster 8; **PKHD1**), hepatic stellate cells (cluster 6; **MYH11**), immune populations (clusters 3, 4, and 9; **PTPRC**, **DOCK2** and **MARCO**), cycling or progenitor-like cells (cluster 7; **TOP2A**), and megakaryocytes (cluster 10; **ITGA2B**).

Based on transcriptional programs, hepatocyte clusters were characterized by the strong expression of genes involved in lipid metabolism and protein secretion, including **ALB**, **APOB**, **ACACA**, **FASN**, and avian-specific **VTG** genes, reflecting their central metabolic and reproductive roles. Endothelial cells displayed heterogeneous endothelial signatures, combining markers associated with sinusoidal endothelial cells (**STAB1**, **STAB2**) and vascular endothelial cells (**VWF**, **KDR**). Immune clusters showed enrichment of canonical leukocyte markers such as **PTPRC**, together with genes involved in immune signaling and antigen presentation, indicating the presence of multiple immune subtypes. Stellate cells were defined by the expression of contractile and extracellular matrix-associated genes, including **MYH11**, **CALD1**, and **COL5A1**, consistent with a perivascular mesenchymal identity. The proliferating cell cluster was characterized by a strong enrichment of cell cycle-related genes (**TOP2A**, **CENPF**, **TPX2**), reflecting an actively dividing population independent of a specific lineage (Supplementary Figure 3). Cholangiocytes were identified by the specific expression of biliary-associated genes such as **PKHD1** and **FGFR2**, supporting their epithelial ductal identity. Finally, megakaryocytes formed a rare but distinct cluster defined by platelet-lineage markers including **ITGA2B** and genes involved in coagulation and cytoskeletal remodeling.

Further details on cluster assignment are provided in the Supplementary Analysis section.

#### Subclustering of major hepatic cell populations reveals intra-lineage heterogeneity

Given their high proportions and transcriptional diversity, hepatocytes (cluster 1 and 5), endothelial cells (cluster 2) and immune cells (cluster 3,4 and 9) were analyzed each one independently to refine cellular subtypes and reduce annotation ambiguity in the context of limited chicken-specific marker genes.

#### Hepatocyte subpopulations (clusters 1 and 5)

To investigate hepatocyte heterogeneity, clusters 1 and 5 were subclustered at a resolution of 0.3 selected based on cluster stability (Supplementary Figure 2B), revealing nine transcriptionally distinct hepatocyte subpopulations (Figure 2A). However, Hep_8 and 9, two very small clusters (combined, 0.1% of hepatocytes) were not further considered due to limited cell numbers (respectively 2 and 89) and weak support from UMAP topology.

**Figure 2.**
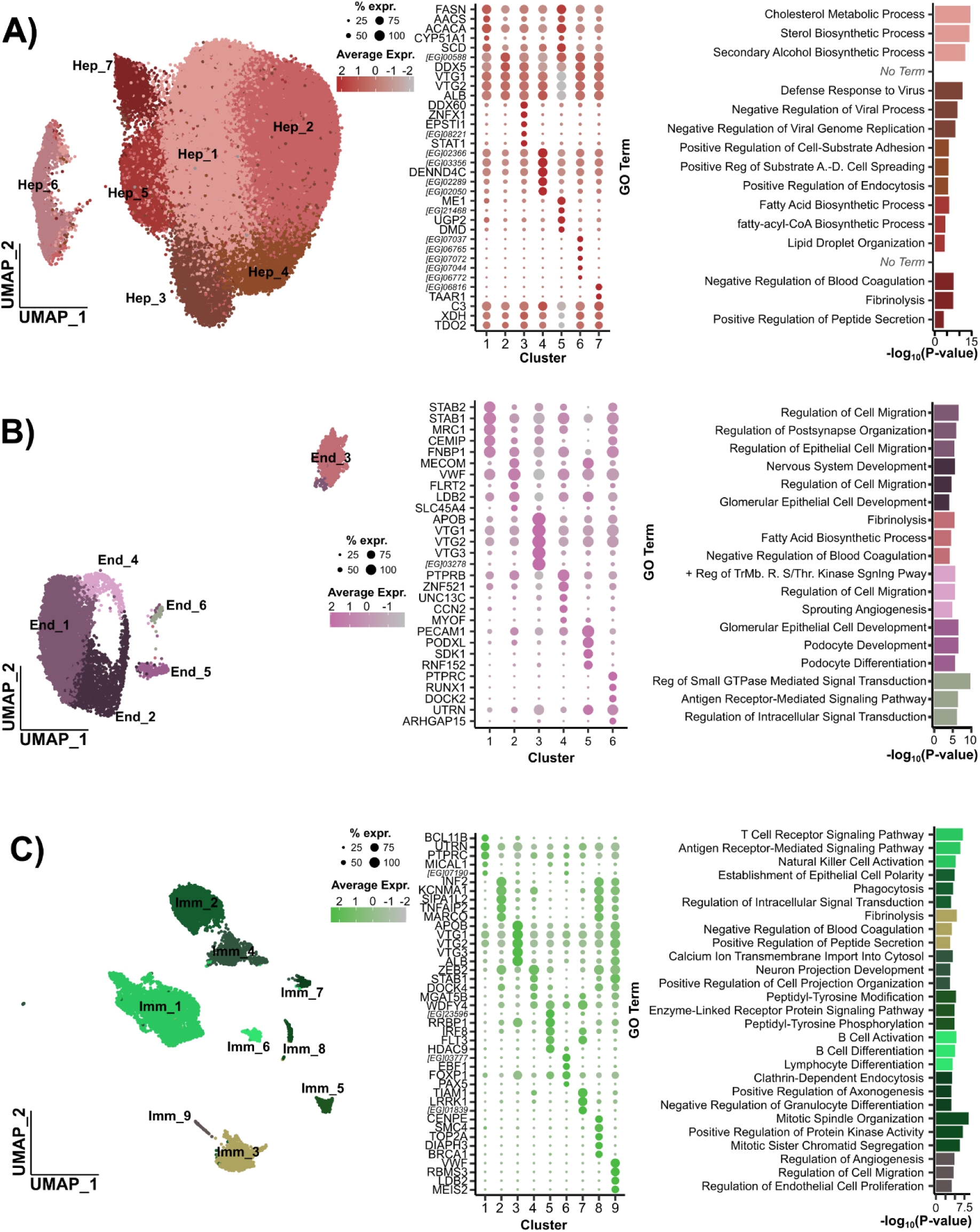
Subclustering reveals transcriptional heterogeneity within major hepatic cell populations. UMAP representation of nuclei subsets with the expression of the top five marker genes per cluster (highest AUC), with dot size representing the proportion of expressing cells and color intensity indicating average expression level. Top three Gene Ontology (GO) biological processes (highest −log10 *p*-value) associated with cluster-specific marker genes. All this information is available for **(A)** Hepatocytes (Hep), **(B)** Endothelial cells (End) and **(C)** Immune (Imm) cells. Expr.: expression; [EG]: ENSGALG000100; A.-D.: Adhesion-Dependent; TrMb. R.: Transmembrane Receptor; S/Thr.: Serine/Threonine.

##### Hep_1 - Lipogenic hepatocytes (n = 34,661 ; 39.0%)

Hep_1 was characterized by enrichment of genes involved in lipid biosynthesis and metabolic processing. Key lipogenic enzymes such as **FASN** (AUC = 0.63; 97.3% pct-in vs 87.6% pct-out), **ACACA** (AUC = 0.61; 97.1% vs 88.6%) and **ACLY** (AUC = 0.60; 76.8% vs 55.5%) indicate an active de novo fatty acid synthesis program. This signature is reinforced by **AACS** (AUC = 0.63; 65.5% vs 39.1%), supporting lipid synthesis from ketone-derived substrates. Genes involved in fatty acid modification (**SCD**, **ELOVL6**) and lipid storage (**PLIN2**, **PNPLA3**) are also enriched, consistent with an active lipogenic program.

GO term analysis highlights strong enrichment for cholesterol and sterol biosynthetic processes as well as broader lipid metabolism involving genes such as **ACLY**, **HMGCS1**, **CYP51A1**, **FDFT1**. Although many of these genes are broadly expressed across the other hepatocytes subclusters, their increased expression in this subcluster reflects a reinforced metabolic state rather than a distinct lineage. Altogether, Hep_1 corresponds to lipogenic hepatocytes with enhanced lipid and cholesterol synthesis, processing and storage capacity.

##### Hep_2 - Basal hepatocytes (n = 32,078 ; 36.1%)

Hep_2 exhibited a transcriptional profile characterized by strong expression of **VTG1**, **VTG2 and VTG3** (AUC ∼0.54; >99% in both cluster and others), which encode avian yolk proteins, together with canonical hepatic genes such as **ALB** (AUC = 0.53; 98.0% vs 99.1%) and **APOB**, reflecting core hepatic synthetic and secretory functions in laying hens rather than a cluster-specific program.

Additional genes, including **SULT6B1L** and **KCNB2**, further supported general metabolic activity. No significant GO term enrichment was detected due to the overrepresentation of VTG genes absent from the database.

Overall, Hep_2 appears to represent a basal hepatocyte population maintaining core hepatic functions in the laying hen liver, while also reflecting the endocrine and metabolic adaptations associated with yolk precursor production.

##### Hep_3 - Immune-responsive hepatocytes (n = 5,795 ; 6.5%)

Hep_3 displayed a distinct transcriptional program consistent with an interferon-responsive / immune-activated hepatocyte state. Key genes such as **DDX60** (AUC = 0.77; 70.1% vs 23.6%), **ZNFX1** (AUC = 0.77; 59.8% vs 8.0%), **EPSTI1** (AUC = 0.71; 51.2% vs 9.8%) and **STAT1** (AUC = 0.71; 55.0% vs 16.5%), indicated activation of innate immune and interferon pathways. GO term enrichment strongly supports this interpretation, highlighting antiviral and interferon-related processes including defense response to virus, antiviral innate immune response, and regulation of interferon-α production. This signature is further reinforced by RNF213 (AUC = 0.69; 59.3% vs 25.7%) and HELZ2 (AUC = 0.61; 35.6% vs 15.5%), together with complement-related genes (C4, C1S), consistent with a stress- and cytokine-responsive state.

##### Hep_4 - Structurally active hepatocytes (n = 5,595 ; 6.3%)

Hep_4 was characterized by genes associated with intracellular trafficking, cytoskeletal organization, and membrane-associated functions, including **DENND4C** (AUC = 0.88; 98.4% vs 71.8%), **PALLD** (AUC = 0.74; 63.4% vs 21.1%), and **LHFPL5** (AUC = 0.68; 45.3% vs 12.0%), although no highly specific marker was identified for this cluster. GO term enrichment highlighted processes related to cell adhesion and tissue organization, including regulation of cell-substrate adhesion and heterotypic cell-cell adhesion. Overall, Hep_4 appears to correspond to structurally hepatocytes with enhanced adhesion, trafficking, and membrane-associated functions rather than a distinct metabolic specialization.

##### Hep_5 - Lipogenic-metabolic hepatocytes (n =4,576 ; 5.2%)

Hep_5 displayed a coherent transcriptional program consistent with a lipogenic (explaining the proximity with Hep1 on the UMAP) and carbohydrate-metabolizing hepatocyte state. GO term enrichment confirmed a strong activation of fatty acid biosynthetic and metabolic processes, as well as fatty-acyl-CoA biosynthesis and lipid droplet organization, driven by key genes such as **FASN**, **ACACA**, **SCD**, **ELOVL6**, **ACSBG2**, **PLIN2** and **PNPLA3**. **ME1** was the strongest marker (*AUC* = 0.80; 81.1% vs 30.6%), supporting NADPH dependent fatty acid synthesis and an anabolic metabolic profile. This interpretation was reinforced by **FASN** (*AUC* = 0.70; 97.6% vs 91.0%), **SCD** (*AUC* = 0.67; 89.4% vs 72.0%) and **ELOVL6** (*AUC* = 0.67; 79.0% vs 53.1%), indicating active lipid synthesis and remodeling. **UGP2** (*AUC* = 0.7; 70.5% vs 34.8%), and **HKDC1** (*AUC* = 0.67; 41.1% vs 7.6%), further suggested coupling between carbohydrate metabolism and lipid biosynthesis.

##### Hep_6 - lncRNA driven hepatocytes (n = 3,381 ; 3.8%)

Hep_6 displayed a distinct transcriptional profile, largely driven by unannotated chicken-specific lncRNA genes (**ENSGALG00010007037**, **ENSGALG00010006765, ENSGALG00010007072 and ENSGALG00010007044)** which are highly enriched in this cluster compared to other hepatocyte subpopulations (*AUC* = 0.7; >36% vs >0.5%). This pattern suggests a coherent and specific transcriptional program rather than minor variation of shared hepatocyte genes. Although functional interpretation is limited by the lack of annotation, the presence of additional markers such as **CACNA1H**, **XDH**, and **AGPAT2** supports a clear hepatocyte identity and suggests a potentially metabolically distinct state. No GO term enrichment was detected, likely due to the predominance of poorly annotated chicken-specific genes, which limits functional annotation in GO databases largely derived from mouse and human references.

##### Hep_7 - Secretory/specialized hepatocytes (n = 2,604 ; 2.9%)

Hep_7 displayed moderate enrichment of plasma protein and coagulation-associated genes, including **FGB**, **FGG**, **PLG**, and **SERPINF2**, with GO term analysis highlighting fibrinolysis and regulation of blood coagulation. Given the relatively broad expression of these markers across hepatocyte populations, Hep_7 likely reflects a continuum of increased coagulation-associated transcriptional activity rather than a discrete hepatocyte subtype.

#### Cross-species comparison of zonation markers

Given this apparent continuum organization, we next assessed whether hepatocyte heterogeneity reflects a canonical mammalian-like metabolic zonation pattern. In mammals, single-cell and spatial transcriptomic studies have revealed a well-defined metabolic zonation of hepatocytes along the porto-central axis. However, a recent cross-species single-cell study (Kaessmann et al.) suggested that such zonation may be reduced or absent in birds. Consistent with this, we examined canonical mammalian zonation markers and observed uniform expression of **GLUL** and **GLDC** across hepatocyte subpopulations, together with weak or negligible expression of **SLC1A2** (Supplementary Figure 4). Together, these findings do not provide evidence for a mammalian-like zonation organization in the laying hen liver.

#### Endothelial subpopulations (Cluster 2)

Within the endothelial cluster (cluster 2), subclustering at a resolution of 0.3, chosen for stability (Supplementary Figure 2C), revealed six transcriptionally different subpopulations (Figure 2B).

##### End_1 - LSEC (Liver Sinusoidal Endothelial Cells) (n =4,709 ; 55.3%)

Based on strong enrichment of canonical scavenger receptors, including **STAB2** (AUC = 0.83; 84.4% vs 40.5%), **STAB1** (AUC = 0.81; 93.6% vs 76.7%) and **MRC1** (AUC = 0.81; 84.9% vs 49.9%), which are associated with endocytosis and clearance of circulating macromolecules, subcluster End_1 was identified as liver sinusoidal endothelial cells (LSECs). Additional markers such as **CEMIP** (AUC = 0.77; 81.0% vs 43.3%), associated with hyaluronan degradation, together with genes associated with cytoskeletal organization and membrane dynamics (**DOCK4**, **FNBP1**, **FCHSD2**) further support an active remodeling phenotype. GO terms enrichment analysis highlighted pathways related to cell migration, cytoskeletal remodeling and VEGF signaling consistent with the dynamic vascular functions of sinusoidal endothelial cells.

##### End_2 - LVEC (Liver Vascular Endothelial Cells) (n =1,645 ; 19.3%)

End_2 subcluster was assigned to vascular endothelial cells, characterized by strong enrichment of **MECOM** (AUC = 0.77; 70.4% vs 26.0%), **VWF** (AUC = 0.73; 92.6% vs 82.6%), **PECAM1** (AUC = 0.64; 55.2% vs 35.5%) and **NR2F2** (AUC = 0.63; 52.8% vs 38.6%) consistent with a non sinusoidal identity involved in vascular integrity and hemostatic function. Gene Ontology analysis highlighted vascular development, endothelial patterning, and regulation of cell migration. Together, these features support annotation of this cluster as a large-vessel/central vein–like endothelial cells distinct from sinusoidal endothelial populations.

##### End_3 - Contaminant - Hepatocyte-like (n =1,377 ; 16.2%)

The strong expression of **APOB**, **ALB**, and vitellogenins (**VTG1/2/3**), together with enrichment of lipid biosynthesis and coagulation related pathways implied an hepatocyte-like transcriptional program. This subcluster likely reflects contaminating hepatocyte-derived or ambient RNA-derived transcriptional signals within the endothelial dataset.

##### End_4 - RLVEC (Remodeling Liver Vascular Endothelial Cells) (n =445 ; 5.2%)

End_4 was characterized by vascular endothelial remodeling features, with enrichment of **PTPRB** (*AUC* = 0.76; 87.0% vs 58.6%), **CCN2** (*AUC* = 0.69; 52% vs 16.5%), **MYOF** (*AUC* = 0.68; 39.1% vs 3.8%) **and ROBO1** (*AUC* = 0.65; 33.7% vs 3.6%). Gene ontology analysis highlighted regulation of cell migration, sprouting angiogenesis, and artery development. Overall, this cluster corresponds to non-sinusoidal endothelial cells involved in vascular remodeling.

##### End_5 - SLVEC (Specialized Liver Vascular Endothelial Cells) (n =247 ; 2.9%)

End_5 corresponded to a specialized non-sinusoidal vascular endothelial subpopulation characterized by enrichment of **PECAM1** (AUC = 0.88; 90.7% vs 37.8%), **PODXL** (AUC = 0.83; 79.8% vs 19.4%), **MECOM** (AUC = 0.80; 80.2% vs 33.2%). Gene Ontology analysis highlighted cell adhesion, endothelial signaling, and regulation of cell migration, together with podocyte-related developmental programs. Overall, this cluster represents a distinct endothelial subpopulation involved in vascular adhesion and tissue architecture remodeling.

##### End_6 - Immune associated cells (n =91 ; 1.1%)

End_6 corresponded to an immune/hematopoietic subpopulation characterized by strong enrichment of **PTPRC** (AUC = 0.80; 61.5% vs 1.2%), **RUNX1** (AUC = 0.73; 47.3% vs 0.9%), DOCK2 (AUC = 0.73; 47.3% vs 1.6%), consistent with leukocyte lineage identity. GO terms supported immune signaling and cytoskeletal regulation.

#### Immunitaire subpopulations (Cluster 3, 4 and 9)

To further resolve immune heterogeneity within the chicken liver, immune cells from clusters 3, 4 and 9 were reanalyzed and subclustered at a resolution of 0.1 (Supplementary Figure 2D), revealing nine transcriptionally distinct immune populations (Figure 2C).

##### Imm_1 - Lymphoïd - T cells (n = 4,893 ; 40.8%)

Imm_1 displayed a canonical T cell transcriptional program characterized by expression of **BCL11B** (AUC = 0.76; 57.1% vs 8.9%) consistent with T-cell lineage commitment. This identity was reinforced by **CD3E** a core component of the T-cell receptor (TCR) complex, and signaling molecules such as **FYN** (AUC = 0.63; 37.0% vs 15.0%) and **PRKCH** (AUC = 0.62; 31.5% vs 9.9%), both involved in TCR-signaling. The enrichment of **TOX2** (AUC = 0.63; 28.9% vs 4.4%) further supports T-cell differentiation while **PTPRC (CD45)** (AUC = 0.74; 69.0% vs 46.2%), **ITGA4** (AUC = 0.62; 33.2% vs 10.8%) and **DOCK 2** indicated lymphocyte activation and migration capacity. These observations were supported by the GO terms enrichment analysis highlighting T cell receptor and antigen receptor–mediated signaling pathways.

##### Imm_2 - Myeloïd - Kupffer-like macrophages (n = 2,819 ; 23.5%)

With strong expression of **MARCO** (AUC = 0.87; 79.6% vs 9.0%) and **MAFB** (AUC = 0.86; 79.9% vs 12.9%), the subcluster Imm_2 identity was consistent with a tissue resident macrophage population with a Kupffer-like identity. This signature was supported by **TNFAIP2** (AUC = 0.89; 83.3% vs 9.4%), indicative of inflammatory activation. Gene Ontology analysis highlighted enrichment of phagocytosis-related processes consistent with active phagocytic activity and tissue debris clearance. Although **PECAM1** was also detected (81.9% vs 26.8%), its co-expression with canonical macrophage markers likely reflects immune–endothelial interactions rather than endothelial contamination.

##### Imm_3 - Contaminant - Hepatocyte-like (n = 2,012 ; 16.8%)

Driven by **APOB** (AUC = 0.99; 99.8% vs 42.4%), **ALB** (AUC = 0.96; 98.7% vs 29.2%), and vitellogenins (VTG1–3), subcluster Imm_3 did not correspond to a true immune population but instead showed a strong hepatocyte-like transcriptional signature. As for endothelial cells, this cluster likely reflects hepatocyte contamination.

##### Imm_4 - Myeloïd - Macrophage/APC (n = 1,226 ; 10.2%)

Imm_4 displayed a myeloid transcriptional program characterized by expression of **ZEB2** (AUC = 0.88; 95.7% vs 46.9%), **STAB1** (AUC = 0.80; 66.2% vs 10.6%), and **CSF1R** (AUC= 0,64; 44.0% vs 18.5%), consistent with a macrophage-lineage identity. This signature was further supported by genes involved in cytoskeletal remodeling and intracellular trafficking, including **DOCK4** (AUC = 0.74; 66.0% vs 24.1%). The expression of **WDFY4** (AUC = 0.71; 63.3% vs 25.3%) and **MGAT5B** indicated the presence of antigen-processing and endosomal functions. Gene Ontology enrichment highlighted receptor tyrosine kinase signaling and phosphorylation-related pathways consistent with an activated myeloid state with antigen-presenting capacity.

##### Imm_5 - Myeloïd - Dendritic cell-like (n = 351 ; 2.9%)

Imm_5 displayed a dendritic cell–like transcriptional program characterized by strong expression of **IRF8** (AUC = 0.86; 79.8% vs 15.8%) and **FLT3** (AUC = 0.85; 72.4% vs 2.5%), together with **WDFY4** (AUC = 0.80; 79.2% vs 27.7%), consistent with an antigen-presenting dendritic-like identity. Gene Ontology analysis highlighted enrichment of receptor-mediated signaling and tyrosine kinase activity consistent with immune activation and environmental sensing.

##### Imm_6 - Lymphoïd - B cells (n = 254 ; 2.1%)

The Imm_6 subcluster exhibited a strong B-cell lineage program characterized by the coordinated expression of key transcriptional regulators, including **EBF1** (AUC = 0.78; 55.9% vs 0.6%), **PAX5** (AUC = 0.70; 39.4% vs 0.2%) and **BCL11A** (AUC = 0.67; 36.6% vs 2.6%). Associated genes linked to signalling such as **MEF2C**, **PLCG2**, and **HDAC9** are also expressed. These genes define a canonical B-cell identity, consistent with B-cell specification, activation, and maintenance. In particular, **PAX5** represents a master regulator of B-cell commitment, while **PLCG2** is a central component of B-cell receptor signaling. Gene Ontology analysis strongly supported this interpretation, revealing significant enrichment of B-cell activation, B-cell differentiation and B-cell receptor signaling pathway. The absence of enrichment for T-cell (e.g., **CD3E**, **BCL11B**) and myeloid markers (e.g., **MARCO**, **CSF1R**) further confirmed a specific B-lineage identity.

##### Imm_7 - Myeloïd - Dendritic cell-like “mature” (n =186 ; 1.6%)

Imm_7 showed a dendritic cell–like transcriptional program characterized by strong expression of key antigen-presenting and differentiation-associated genes, including **FLT3** (AUC = 0.87; 75.3% vs 3.4%), **IRF8** (AUC = 0.76; 63.4% vs 17.0%), **WDFY4** (AUC = 0.88; 87.6% vs 28.3%), and **CADM1** (AUC = 0.84; 74.7% vs 15.1%), consistent with a dendritic cell differentiation and antigen presentation identity. Additional markers such as **TIAM1** (AUC = 0.92; 87.6% in-cluster vs 8.3% out-of-cluster), **LRRK1** (AUC = 0.91; 83.9% vs 8.5%), **PLCB2** (AUC = 0.79; 69% vs 19%), and **PTK2** (AUC = 0.76; 65% vs 24%) indicated strong cytoskeletal remodeling and intracellular signaling activity, suggesting an antigen-sampling and migratory immune phenotype. Gene Ontology analysis revealed enrichment of clathrin-dependent endocytosis (**ITSN2** and **FCHSD2**), cytoskeletal remodeling and regulation of differentiation (**RUNX1** and **INPP5D**), supporting a highly active migratory dendritic cell state. Overall, this cluster represents a mature dendritic cell–like population.

##### Imm_8 - Myeloïd - Proliferating Kupffer-like macrophages (n =167 ; 1.4%)

Showing a strong cell-cycle program, subcluster Imm_8 was considered as proliferating macrophages. This proliferative state was driven by **CENPE** (AUC = 0.82; 64.1% vs 0.4%), **TOP2A** (AUC = 0.80; 61.1% vs 0.5%), **SMC4** (AUC = 0.81; 71.9% vs 9.7%), **DIAPH3** (AUC = 0.80; 61.1% vs 0.6%) and **BRCA1** (AUC = 0.79; 58.7% vs 1.0%), confirming active progression through mitosis (S/G2-M phases). Importantly, this signature was coexisting with a macrophage program marked by **MARCO**, **MAFB**, **TNFAIP2**, **INF2**, and **PLXNB2**, indicating a Kupffer-like identity. Gene Ontology enrichment confirmed mitotic activity together with kinase signaling regulation.

##### Imm_9 - Endothelial-like (n =71 ; 0.6%)

Cluster 9 showed a clear endothelial contamination signature rather than an immune profile, driven by strong expression of **VWF**, **KDR** and **PLXND1**, together with **STAB1**, **STAB2** and **MRC1**.

#### Lineage-specific subclustering improves the resolution of hepatic cell populations

As an alternative to lineage-specific subclustering, the complete dataset was also clustered at a higher resolution (resolution = 1.0 instead of 0.1; Supplementary Figure 2A). The resulting high-resolution global clustering identified 23 clusters (Supplementary Figure 5A), comprising nine hepatocyte, two endothelial and eight immune populations, together with the same four stable macro-populations (cholangiocytes, stellate cells, megakaryocytes and proliferating cells). We then compared this strategy with lineage-specific subclustering, in which hepatocyte, endothelial and immune compartments were independently reclustered before being merged into a single dataset. This approach resulted in 28 populations (Supplementary Figure 5B), including nine hepatocyte, six endothelial and nine immune subclusters, together with the same four stable macro-populations.

Although both strategies recovered the same major hepatic compartments, they differed in their ability to resolve intra-lineage heterogeneity. High-resolution global clustering did not always separate closely related cell states, particularly within hepatocytes, where several global clusters overlapped with multiple subclusters obtained by lineage-specific analysis (Supplementary Figure 5C). In contrast, lineage-specific subclustering improved the identification of endothelial and immune populations, resolving six endothelial subpopulations and distinct myeloid and lymphoid states. Overall, these results indicate that lineage-specific subclustering provides a finer resolution of cellular diversity within major hepatic compartments.

## IMPACT OF AN ENRICHED ANNOTATION ON SINGLE-NUCLEUS RNA-SEQ ANALYSIS

To assess the influence of gene annotation on the cell population identification and downstream biological interpretation, we repeated clustering and marker gene identification using an enriched annotation combining RefSeq, Ensembl, and four additional transcriptomic resources (see Materials and Methods). Results were then compared with those obtained using the standard Ensembl reference annotation described above.

### General statistics

Using the same workflow as for previous analysis, after quality-control filtering based on the distributions of the number of genes and transcripts detected (Supplementary Table 5), 118,780 nuclei were retained for downstream analyses (+1.3% compared to analysis with Ensembl reference annotation only). The median number of nuclei recovered per sample was 9509 (+1.1%), with the same distribution across samples as previously observed. Sequencing depth was overall homogeneous across nuclei and across samples, with a median of approximately 7,548 (+8.3%) reads per nucleus. The median number of genes detected per nucleus was 941, (+9.2%) showing limited inter-sample variation. Likewise, the median number of transcripts per nucleus was 1,785 (+14.3%) across samples, indicating robust and consistent transcript capture throughout the dataset.

### A limited impact on global clustering

At the macro-cluster level, both annotation strategies produced highly similar cellular organizations. Using the enriched annotation, integrated single-nucleus RNA-seq analysis resolved eleven transcriptionally distinct hepatic cell populations in the chicken liver (Figure 3A). Hepatocytes represented the dominant population (ClusterEA_1; 71.9% of nuclei), whereas endothelial, immune, stellate, cholangiocyte, proliferating, and megakaryocyte populations formed smaller but clearly separated clusters as observed with the Ensembl annotation. The relative proportions of these major cell populations remained consistent across the eight individuals, supporting the robustness and reproducibility of the clustering strategy (Figure 3B).

**Figure 3.**
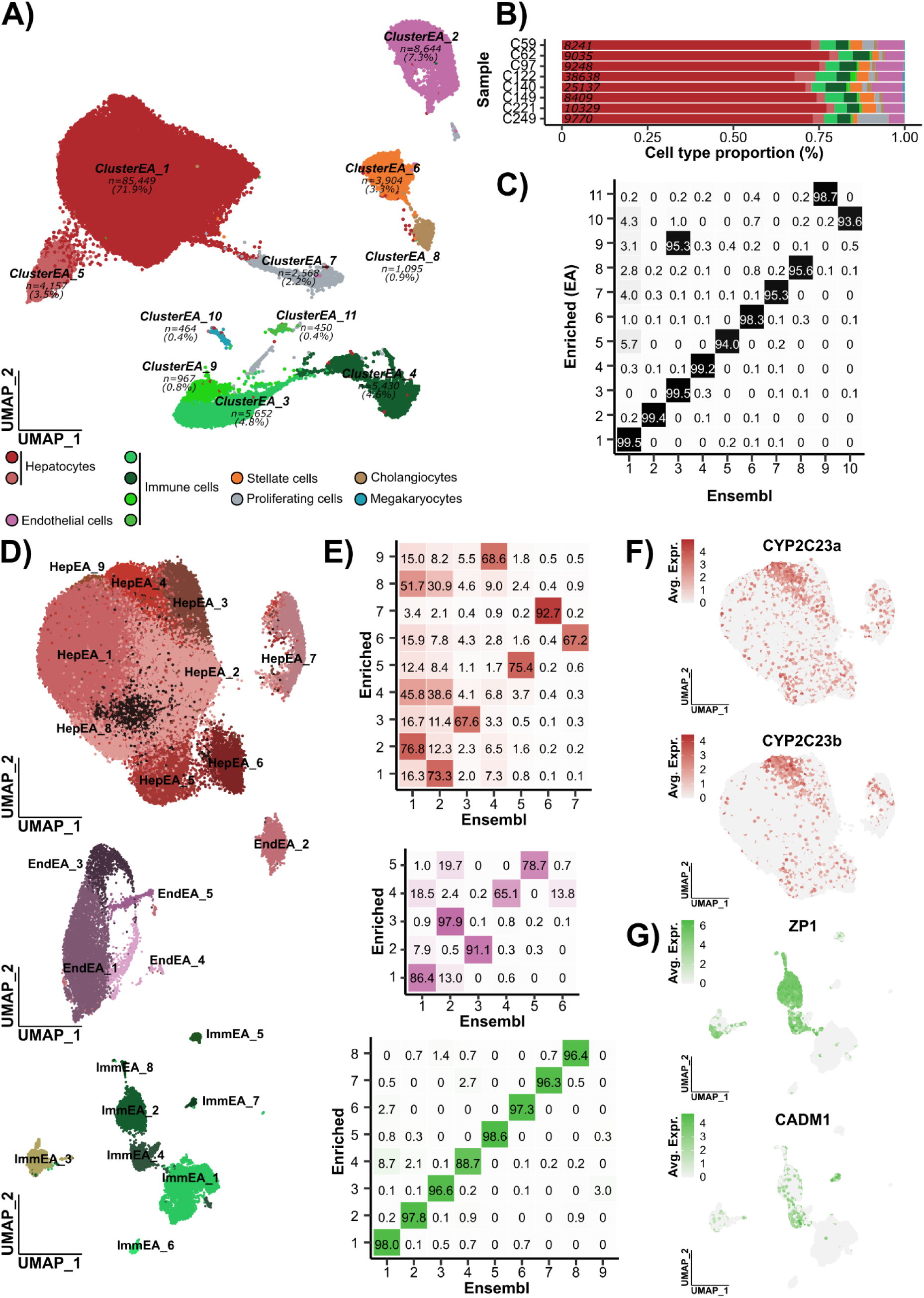
Enriched annotation impact on chicken liver cell-type identification and annotation. **(A)** UMAP of integrated single-nucleus RNA-seq data using the enriched annotation highlighting eleven transcriptionally distinct clusters (indicating in italics). (n=) indicates the number of cells associated with the cluster and the corresponding percentage. Cluster biological identities were assigned based on marker genes and correspondence with previous assignment shown in panel C). **(B)** Proportion of each major cell type across the eight individual samples; numbers in italics indicate the total number of nuclei per sample. Colors correspond to the cell-type identities shown in panel (A). **(C)** Concordance matrix comparing clusters identified using the enriched annotation workflow (“Enriched”) with previously defined reference annotations (“Ensembl”). Values represent the percentage overlap considering the “Enriched” clusters as reference. **(D)** UMAP representation and **(E)** correspondence matrices of subclustering analyses for hepatocytes (top - red shade), endothelial cells (middle - purple shade), and immune cells (bottom - green shade) obtained following enriched annotation. Expression of marker genes **(F)** *CYP2C23a* and *CYP2C23b* across hepatocyte subclusters and **(G)** *ZP1* and *CADM1* across immune subclusters. Avg. Expr.: Average Expression.

To evaluate the impact of the enriched annotation on cell-type assignment, we compared the resulting clusters with those previously obtained using the Ensembl-based workflow. The concordance matrix revealed an almost perfect correspondence between annotations, with overlap values generally exceeding 95% (Figure 3C) and supported by a high ARI index (0.97). Canonical lineage markers and top-ranked lineage markers remained highly consistent between analyses (Table 1) with around 40 genes conserved over the 50th considered using each annotation. Differences between annotation strategies primarily involved genes absent from the standard Ensembl annotation, alternative gene models resulting from transcript fusion or splitting events, and several lncRNAs identified only in the enriched annotation. For example, **TAT**, a canonical hepatocyte-associated gene, was absent from the standard Ensembl annotation but was recovered using the enriched annotation (RefSeq gene model origin) and contributed to hepatocyte specific signatures.

**Table 1.**
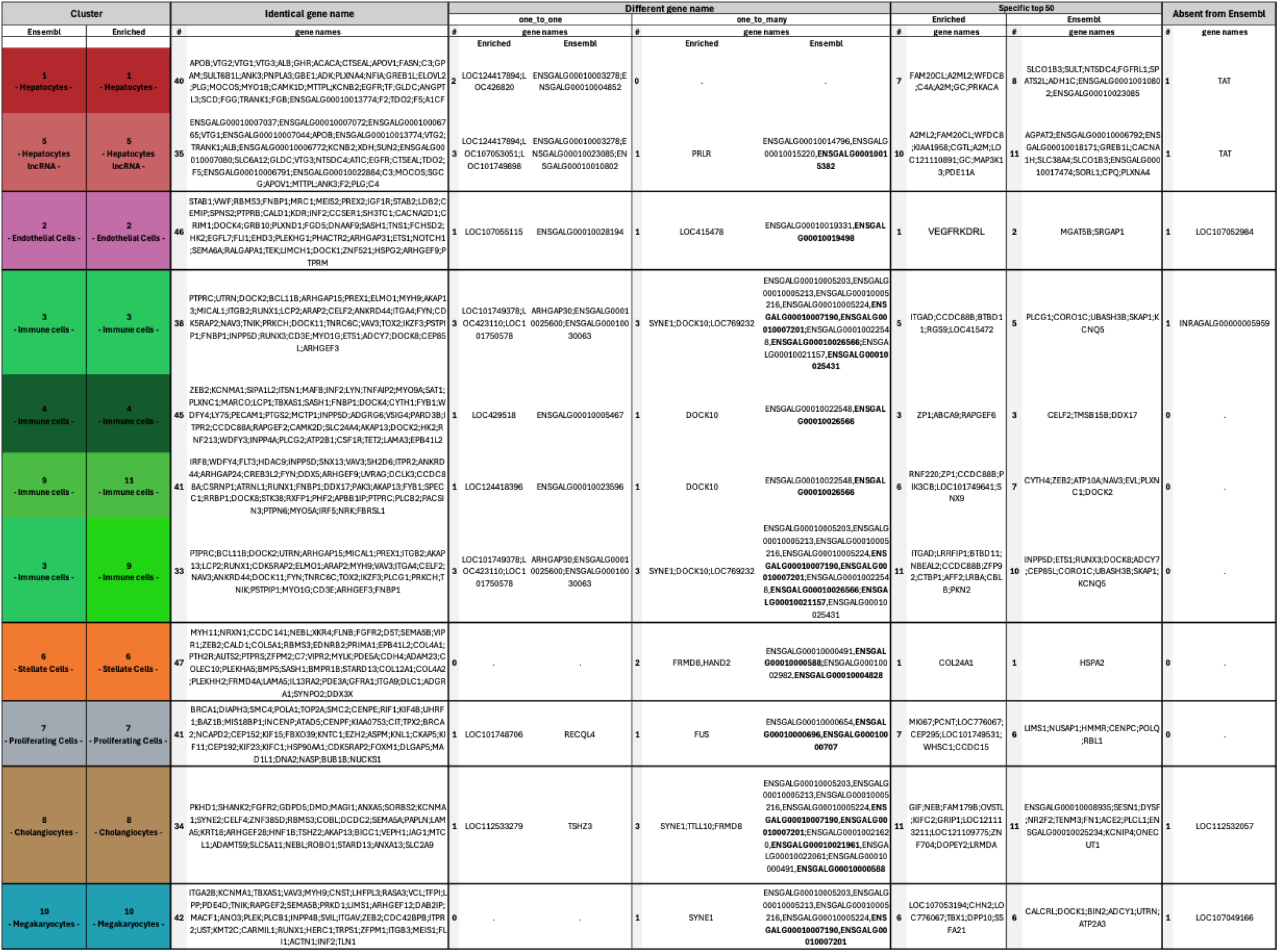
Comparative analysis of liver cell cluster annotations obtained using Ensembl and enriched annotations. The “Identical gene name” section lists conserved marker genes detected with the same gene symbols in both annotations. The “Different gene name” section highlights markers corresponding to equivalent genes but annotated under different identifiers, including one-to-one and one-to-many relationships between annotations. The “Specific top 50” columns report marker genes uniquely identified among the top 50 cluster markers in either the enriched or Ensembl annotation. The “Absent from Ensembl” column indicates genes detected exclusively with the enriched annotation and lacking corresponding annotation in Ensembl. Numbers (#) indicate the count of genes in each category.

### Impact on cluster-specific biology

We next investigated whether the enriched annotation increased resolution within major cellular compartments by performing dedicated subclustering analyses of hepatocytes, endothelial cells, and immune cells (Figure 3D). Hepatocyte analysis identified nine subclusters exhibiting heterogeneous transcriptional programs, whereas endothelial and immune populations segregated into five and eight subclusters, respectively. Comparison with the Ensembl-derived subclusters demonstrated variable levels of correspondence depending on the cell compartment (Figure 3E). Immune populations showed the strongest conservation between annotations, with nearly all subclusters displaying >95% overlap (ARI = 0.93). In contrast, hepatocyte (ARI = 0.37) and endothelial (ARI = 0.69) populations exhibited more substantial redistributions, indicating that gene models from Enriched annotation affected the resolution of these transcriptionally complex populations. Several hepatocyte subclusters exhibited mixed correspondence patterns rather than one-to-one matches, consistent with the notion that hepatocyte transcriptional organization reflects a continuum of cellular states rather than sharply separated populations.

Importantly, the enriched annotation enabled the detection and characterization of biologically informative marker genes that were poorly captured or absent using the Ensembl annotation. Within hepatocytes, CYP2C23a and CYP2C23b showed restricted expression patterns across the HepEA_4 subcluster (Figure 3F), supporting the existence of functionally specialized hepatocytes. Similarly, in immune populations, ZP1 and CADM1 exhibited specific expression profiles (Figure 3G), allowing a clearer distinction between myeloid- and lymphoid-associated populations and illustrating the increased sensitivity provided by the enriched annotation. In Kupffer-like macrophages, the enriched annotation refined the transcriptional signature by revealing additional markers, including ZP1, while preserving canonical macrophage-associated genes such as MARCO and TNFAIP2. Collectively, these results demonstrate that the enriched annotation has limited impact on global liver cell-type composition but substantially enhances the resolution and interpretability of intra-lineage heterogeneity in chicken liver single-nucleus transcriptomic data.

## Discussion

The present study provides a single-nucleus transcriptomic atlas of the adult laying hen liver, offering a comprehensive view of its cellular composition and transcriptional landscape. Our analysis identified the major hepatic cell populations, including hepatocytes, endothelial cells, cholangiocytes, stellate cells, and diverse immune cell types, highlighting the complexity of the chicken liver. The major hepatic cell populations identified here are consistent with those reported in mammalian liver single-cell atlases, supporting the robustness of our cell type annotation and the evolutionary conservation of the liver cellular architecture across vertebrates.This overall conservation provides additional confidence in the biological relevance of our findings while establishing a species-specific reference for the adult laying hen liver.

To ensure robust identification of hepatic cell populations at the global level, a relatively low clustering resolution was applied. This choice was motivated by the risk that increasing global clustering resolution may lead to sub- or over-optimal fragmentation of cell populations displaying different degrees of specialization. This could lead to the emergence of transcriptionally unstable clusters that are difficult to interpret or biologically meaningless. Following global annotation of major cell types, the three most abundant and transcriptionally diverse compartments (hepatocytes, endothelial cells, immune cells) were independently re-analyzed using sub-clustering approaches. This strategy enabled the resolution of intra-lineage heterogeneity while maintaining stable global cell type definitions.

Hepatocytes represented the dominant cell population (∼70%) and exhibited substantial transcriptional heterogeneity. Several hepatocyte subpopulations were enriched for lipid metabolism pathways, consistent with the central role of the avian liver in yolk precursor synthesis. In particular, the strong expression of the three vitellogenin genes (VTG1–VTG3) together with apolipoprotein B (APOB) reflects a specialized hepatic program dedicated to lipid transport and lipoprotein assembly during egg production. This transcriptional program is driven by estrogen signaling and has been extensively characterized as a coordinated induction of yolk precursor genes during the laying cycle, involving profound metabolic reprogramming of hepatocytes (43,44). This estrogen-dependent specialization underlies the physiological plasticity of the liver in laying hens and contributes to the marked transcriptional diversity observed among hepatocyte subpopulations in our dataset.

Subclustering analysis revealed that hepatocytes were organized along a continuum rather than as strictly discrete populations. A basal hepatocyte population expressing canonical liver genes occupied a central position within the continuum and was connected to more specialized programs, including lipogenic, secretory-associated, interferon-responsive, and structural activity states. This organization suggests graded functional specialization rather than strict compartmentalization. In contrast to mammalian livers, where hepatocyte zonation along the porto-central axis represents a major organizational feature, no clear evidence of such zonation was observed in the chicken liver, consistent with previous observations in avian species (22). However, this interpretation should remain cautious given the absence of spatial resolution information inherent to single-nucleus transcriptomic approaches. Altogether, these observations suggest that hepatocyte diversification in the laying hen may be driven more strongly by physiological and reproductive specialization than by the spatial metabolic gradients classically described in mammals.

Endothelial cells displayed substantial transcriptional heterogeneity. In the present dataset, this diversity was organized into two major transcriptional compartments corresponding to sinusoidal (STAB1, STAB2, MRC1) and vascular (VWF, NR2F2, MECOM) identities. Within these compartments, cells were further distinguished by functional transcriptional programs associated with scavenging activity in sinusoidal endothelial cells and with vascular identity, angiogenic remodeling, or endothelial adhesion in vascular endothelial cells. By contrast to mammalian livers, where endothelial heterogeneity is tightly linked to porto-central vascular organization and the specialized function of liver sinusoidal endothelial cells (45), no clear evidence of spatially organized transcriptional gradients was observed in the chicken liver. Although this interpretation remains limited by the lack of spatial transcriptomic information, these observations suggest that endothelial diversity in laying hens is primarily structured by functional compartmentalization rather than by canonical mammalian-like vascular zonation.

Our analysis also revealed a transcriptionally active immune compartment comprising myeloid and lymphoid populations. Cell-type annotation was guided by conserved lineage-defining gene expression programs across vertebrates and supported by established avian and mammalian literature, enabling robust identification of major immune populations. This is consistent with the known immune architecture of the liver as a central immunological organ (46). Putative Kupffer-like macrophages were characterized by expression of MARCO and MAFB, consistent with a tissue-resident macrophage identity involved in phagocytic and scavenging functions. In parallel, lymphoid populations were identified based on canonical lineage markers, including CD3D and BCL11B for T cells, and PAX5 and CD79A for B cells, reflecting conserved adaptive immune compartments in the chicken liver. Functional enrichment analyses further highlighted active cytokine signaling, T-cell receptor signaling, and MAPK pathway activity, consistent with an immunologically responsive hepatic environment. Within the myeloid compartment, the presence of transcriptionally distinct states with antigen-presenting and activated features indicates functional diversification beyond a single homogeneous macrophage population. Overall, these results support a conserved but functionally active immune landscape in the chicken liver, integrating innate and adaptive immune programs within the hepatic microenvironment.

Comparison with other single cell studies in chicken reveals interesting discrepancies in liver cellular composition across datasets. In particular, recent large-scale initiatives such as ChickenGTEx have provided a valuable multi-tissue single-cell resource in chicken, including liver (16), where 13 cell types were identified from approximately 14,000 cells, In this dataset, a relatively high proportion of Kupffer cells was reported compared to parenchymal populations. These differences likely reflect a combination of biological samples and methodological factors. Among these, dissociation-induced biases inherent to single-cell RNA sequencing workflow together with the known sensitivity of hepatocytes to enzymatic and mechanical processing, which may contribute to their selective underrepresentation (47). In contrast, single-nucleus RNA sequencing improves the recovery of large and fragile parenchymal cells resulting in a more balanced representation of hepatic cell types (48). Furthermore, differences in clustering and data integration strategies across studies may influence the resolution of tissue-specific populations. In our dataset, this effect is illustrated by non-immune contaminants (Imm_3 and Imm_9) which were initially co-captured within low-resolution immune clusters and subsequently resolved through higher-resolution subclustering.

Importantly, similar overall hepatic cellular architectures have also been reported in other avian single cell studies, including embryonic chicken liver atlases (23) and duck liver single-cell datasets (49), despite differences in developmental stage and experimental context. These studies consistently identified the major hepatic compartments, supporting a conserved organization of the avian liver across species and conditions. In contrast to these datasets, the present study focuses on hens sampled at the end of the commercial laying period (∼90 weeks of age), corresponding to a late reproductive stage characterized by sustained egg production and prolonged exposure to estrogen-driven hepatic metabolism. At this stage, the liver operates under sustained metabolic and hormonal demand, which may contribute to long-term adaptive remodeling of hepatic cell states. Although no age-matched direct comparison is available, the transcriptional heterogeneity observed here likely reflects a combination of reproductive status, prolonged physiological activity, and age-associated remodeling of hepatic function. This is notably illustrated by the expression pattern of vitellogenin (VTG) genes, which are virtually absent from the embryonic chicken liver atlas (23) and restricted to a limited hepatocyte subpopulations in the duck liver dataset (49), whereas they are broadly expressed across multiple hepatocyte subclusters in our study. This expanded representation of VTG-expressing hepatocytes likely contributes to the distinct hepatocyte subpopulation structure observed here and underscores the influence of physiological context on avian liver cellular organization.

Another key feature of our study was the use of an enriched annotation for the GRCg7b chicken genome (27,28) which incorporated numerous previously unannotated lncRNA genes and improved the transcript structures of several protein-coding gene models compared with the reference Ensembl annotation. While global cell type composition remained highly consistent between annotation strategies, as supported by strong concordance and high ARI values, the enriched annotation improved the resolution at the subcluster-level, particularly within hepatocyte and immune populations. In practice, the main contribution of the enriched annotation was not the emergence of entirely novel lncRNA-defined cell populations, but rather improved transcript detection and read assignment for biologically relevant genes that were incompletely represented in the standard annotation This was notably illustrated by CYP2C23A and CYP2C23B which were not detected in the Ensembl annotation due to a fusion of these gene models and allowed the detection of previously unidentified hepatocytes subtypes. In the laying hen liver, these genes are known to be estrogen-responsive and expressed in the liver during egg production supporting the interpretation that the refined annotation enabled a more accurate characterization of hepatocyte functional heterogeneity (50).

Interestingly, although lncRNAs were generally not major drivers of global cell-type discrimination, one hepatocyte subpopulation displayed coordinated high expression of several lncRNA genes located within the same chromosomal region. This pattern suggests the presence of localized transcriptional or chromatin-associated regulatory activity rather than isolated stochastic expression events. However, the functional significance of this signal remains unclear and will require further investigation using complementary approaches such as integrative analyses and targeted functional studies.

In contrast to the Ensembl annotation, the enriched annotation also enabled the identification of ZP1 expression within Kupffer-like cells. ZP1 encodes a component of the perivitelline membrane and is classically associated with oocyte biology (51), although is also known to be estrogen-responsive and expressed in the liver during egg production. We observed that ZP1 expression was predominantly localized to Kupffer-like cell clusters rather than hepatocytes. Interestingly, exploration of the avian liver dataset available through the single-cell resource (https://apps.kaessmannlab.org/liver_app/) generated by Kaessmann and al., revealed a similar expression pattern, supporting the reproducibility of this observation. To our knowledge, this represents one of the first descriptions of this specific cellular localization pattern in the adult laying hen liver even if this gene is considered as expressed globally in the liver. This underscores the influence of comprehensive genome annotation in single-nucleus transcriptomic analyses, particularly in non-model organisms where incomplete gene models may obscure biologically relevant expression patterns.

Looking forward, the dataset generated here provides a valuable resource to the community working on the liver and cell specificity of this organ. This resource may be useful for deconvolution of the numerous bulk RNA-seq data from chicken livers available on the public database. This is allowing the inference of cell-type proportions in larger cohorts or under different physiological conditions, such as different stages of the laying cycle. Integration with spatial transcriptomics or functional assays could further refine our understanding of cellular localization and metabolic specialization.

## Supporting information

SupTable1

SupTable2

SupTable3

SupTable4

SupTable5

Supplementary_Analyses

## Acknowledgements

The samples used in this study were generated as part of the European Union’s Horizon 2020 GEroNIMO project (grant agreement N°101000236), and the French National Research Agency (ANR) CE20 project EFFICACE. The authors wish to thank the Novogen breeding company for producing and rearing the animals from which the samples were collected.

## Author contributions

LL and SL conceived and designed the study and secured the funding. LL, BL and CA performed sampling and laboratory work. LL and FD performed bioinformatic analysis. LL and FD analysed the results, drafted the manuscript, and managed the project. All authors read and approved the final version of the manuscript.

## Funding

This study received financial support from the “Genetic Animal” department of INRAE.

## Data reporting

The raw data (FASTQ files) from the snRNA-seq were uploaded to the European Nucleotide Archive (ENA) database under accession number PRJEB115271.

## Ethics approval

The experiments were performed in accordance with animal welfare standards and were approved by the ethics committee for animal experimentation (APAFIS #34628-2022011214434077 v3), in compliance with French and European legislation.

## Supplementary Figures

**Supplementary Figure 1.**
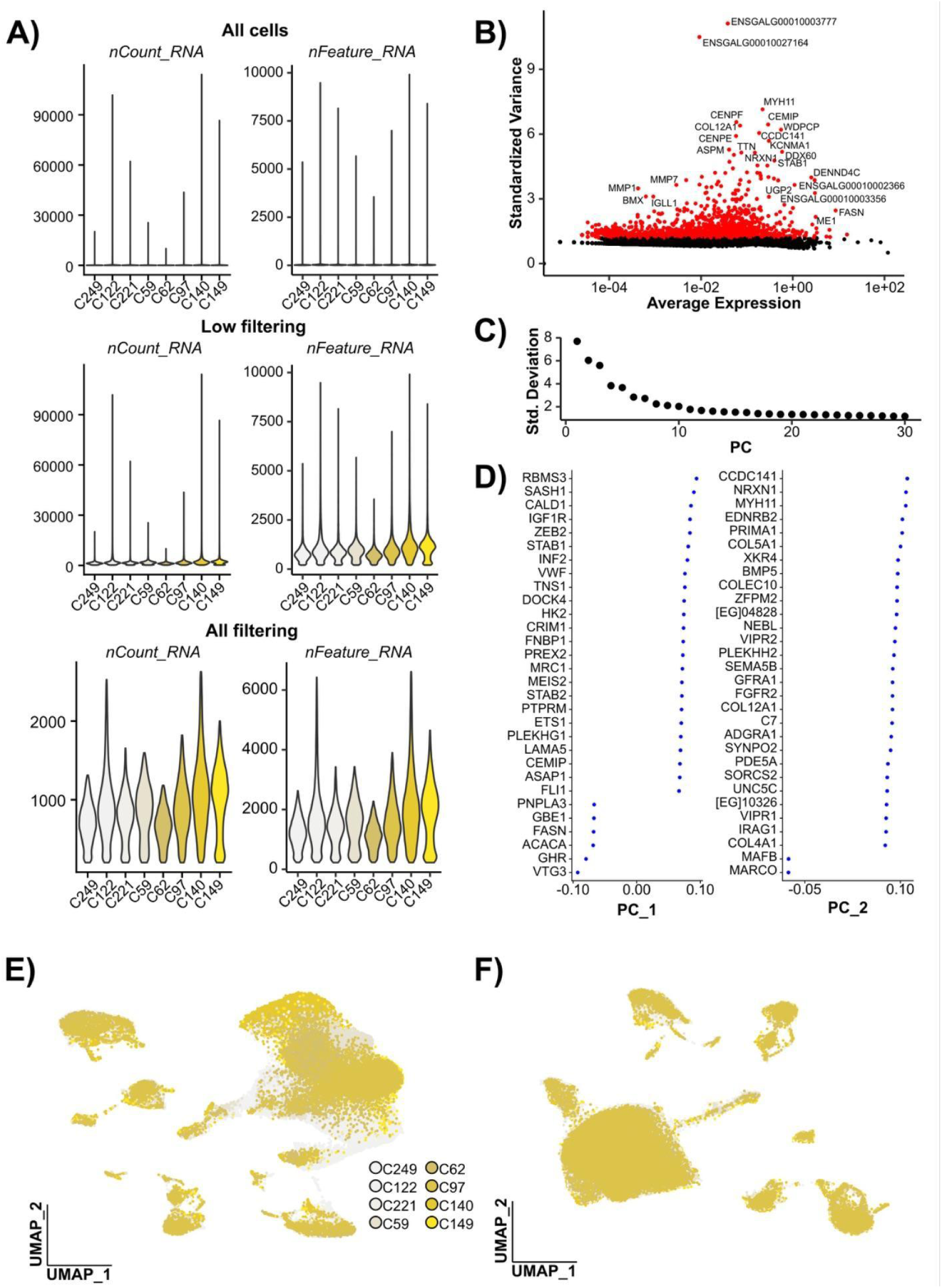
Data preprocessing and dimensionality reduction of single-nucleus RNA-seq data. **(A)** Distribution of sequencing metrics across samples, including number of UMIs (nFeature_RNA) and detected genes (nCount_RNA), shown for all cells (top), after low-stringency filtering (middle), and after final filtering (bottom). **(B)** Identification of highly variable genes based on standardized variance. Genes selected for downstream analyses are highlighted in red. **(C)** Elbow plot displaying the standard deviation explained by each principal component (PC). **(D)** Top genes contributing to the first two principal components (PC1 and PC2), ranked by their loading values. UMAP representation colored by sample origin, illustrating the distribution of nuclei **(E)** prior to integration UMAP and **(F)** after data integration (harmony).

**Supplementary Figure 2.**
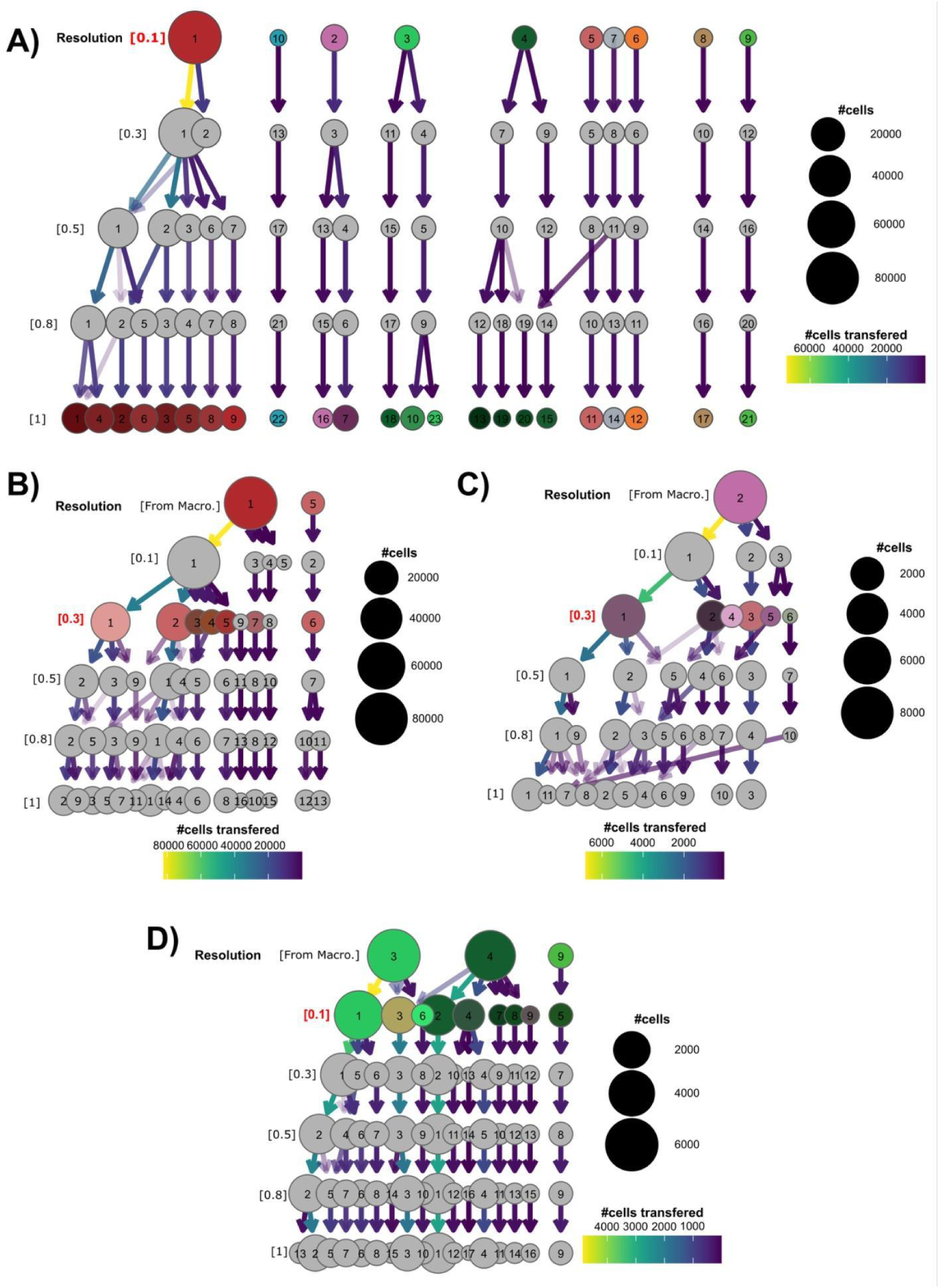
Hierarchical clustering structure across resolutions and lineage-specific subclustering. **(A)** Global clustering tree showing the relationships between clusters across increasing resolutions (from 0.1 to 1). **(B–D)** Lineage-specific clustering trees derived from major cell populations illustrating subclustering trajectories across increasing resolutions for **(B)** hepatocyte lineage, **(C)** endothelial lineage, and **(D)** immune lineage. Across all panels, each node represents a cluster, with node size proportional to the number of cells. Arrows indicate the transition of cells between clusters across resolutions, with color intensity reflecting the proportion of cells transferred. Resolutions highlighted in red correspond to thus kept for the biological analysis.

**Supplementary Figure 3.**
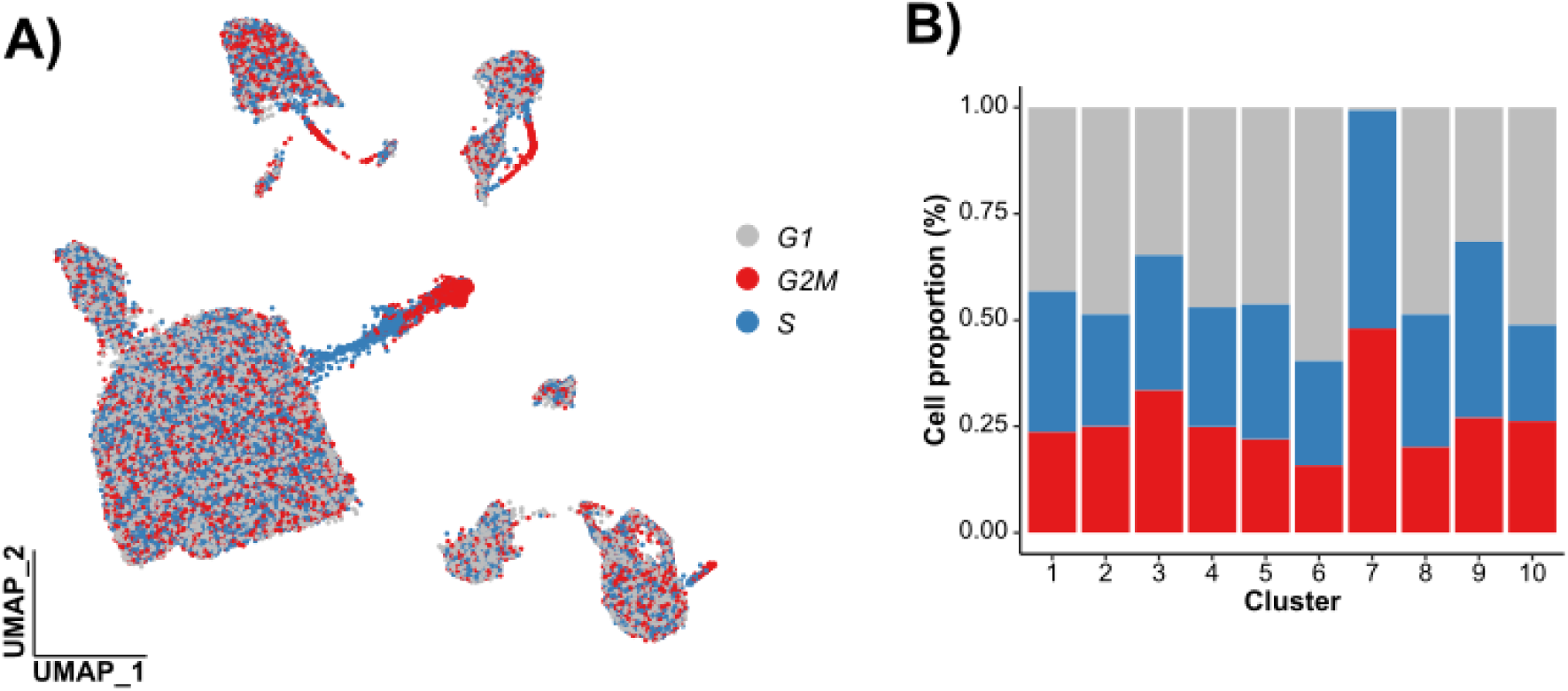
Cell-cycle distribution across cell macroclusters through Ensembl annotation. **(A)** UMAP representation colored according to inferred cell-cycle phase: G1 (grey), S (blue), and G2/M (red). **(B)** Proportion of cells assigned to each cell-cycle phase across the 10 identified clusters.

**Supplementary Figure 4.**
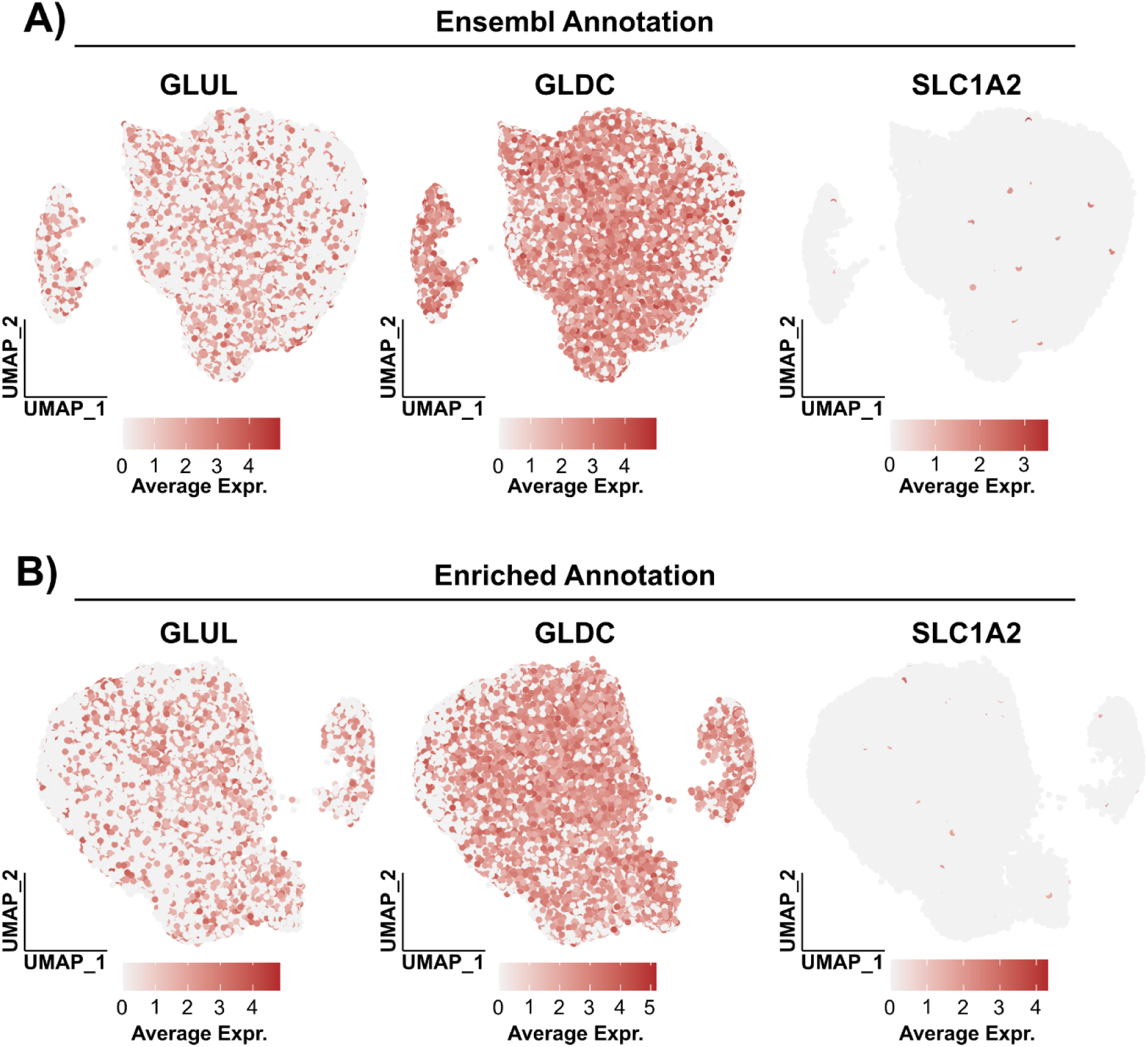
Expression of GLUL, GLDC and SLC1A2 mammalian hepatic zonation marker genes in hepatocyte subclustering analyses according to the **(A)** Ensembl (Clusters 1 and 5) or **(B)** enriched (Clusters EA_1 and EA_5) annotation. Average Expr.: average expression level.

**Supplementary Figure 5.**
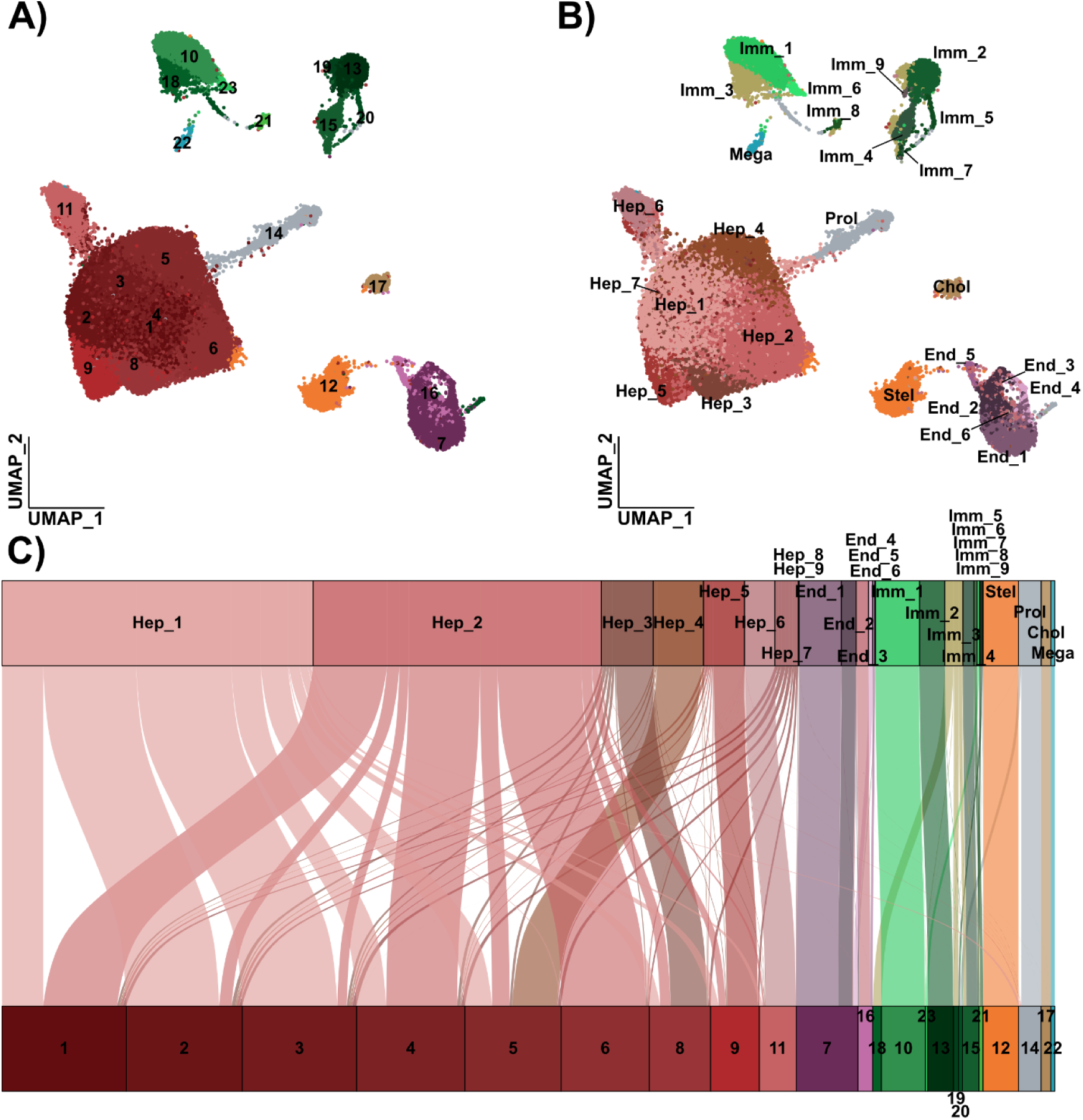
Comparison between lineage-specific subclustering and global clustering approaches. UMAP representation of the dataset colored according to (**A)** clusters identified using a global approach (resolution = 1) or **(B)** concatenated micro-clusters obtained from lineage-specific subclustering analyses, including hepatocyte (Hep), endothelial (Endo), immune (Immu), and minor cell populations. **(C)** Alluvial plot showing the correspondence between lineage-specific micro-clusters (top) and global clustering (bottom). Stream widths are proportional to the number of cells. Colors in **(A)** reflect the combination of major cell population identity and their respective subclusters. In **(B)**, clusters are colored based on their assigned major cell population, with derived shades used to distinguish subclusters when applicable.

## Supplementary Tables

**Sup. Table 1 –** Per-sample and overall statistics of single-nucleus RNA-seq analysis across the successive filtering steps, from raw nuclei detection to the final filtered dataset for the Ensembl annotation.

**Sup. Table 2** – List of marker genes used for cell-type annotation and the corresponding literature references supporting their identification.

**Sup. Table 3** – Gene Ontology (GO) term enrichment analysis associated with hepatocyte, endothelial, and immune cell subclusters.

**Sup. Table 4 –** Summary statistics of the top 50 marker genes used for Gene Ontology (GO) enrichment analyses across hepatocyte, endothelial, and immune cell subclusters.

**Sup. Table 5** – Per-sample and overall statistics of single-nucleus RNA-seq analysis across the successive filtering steps, from raw nuclei detection to the final filtered dataset for the Enriched annotation.

## Notes

### Competing Interest Statement

The authors have declared no competing interest.

