## Supplementary_Analyses for "A single-nucleus atlas of the adult laying hen liver reveals metabolic specialization and improved cellular resolution through enhanced genome annotation"

### Supplementary analysis

##### Annotation of Major Hepatic Cell Populations

###### Hepatocyte populations (Clusters 1 and 5 ; n = 84,719 and 4,059 ; 72.2% and 3.5%)

Cluster 1 was annotated as hepatocytes based on the strong enrichment of genes associated with hepatic metabolic, lipid metabolic, and secretory functions. Among the most discriminant markers, **ALB**, **ACACA**, and **FASN** showed high classification performance (AUC = 0.82, 0.79, and 0.79, respectively), with a large proportion of expressing nuclei within this cluster (>90%) compared to other clusters (<50%). These genes are central to fatty acid synthesis and triglyceride production, which are key metabolic functions of hepatocytes. In addition, this cluster displayed strong enrichment of genes involved in lipoprotein assembly and lipid transport, such as **APOB** (AUC = 0.85) and **APOV1** (AUC = 0.79).

Consistent with the avian model, the vitellogenin genes **VTG1**, **VTG2**, and **VTG3** were also highly enriched (AUC > 0.90), with expression largely restricted to this cluster. These genes are involved in the hepatic synthesis and secretion of yolk precursors and lipid-rich particles, a process strongly stimulated by estrogen during the laying period. Their coordinated expression highlights the specialized role of hepatocytes in supporting oocyte development through the production and transport of vitellogenins and apolipoproteins.

Additional markers further supported hepatocyte-specific functions. **PNPLA3** (AUC = 0.77) was enriched (83% vs 35% of cells), consistent with its role in lipid remodeling and storage. The expression of **GHR** (AUC = 0.81) further supports the endocrine responsiveness of these cells, particularly to growth hormone signaling.

**C3** (AUC = 0.78), a key component of the complement system, showed higher expression in hepatocytes (91% vs 42%), in line with the liver being a major source of circulating complement proteins.

These markers were also detected in cluster 5, although with generally lower AUC values (>0.65) likely reflecting the influence of cluster 1 in the differential comparison (one cluster versus all others). This explains the close proximity of clusters 1 and 5 in the UMAP and supports their shared annotation as hepatocyte populations. However, cluster 5 exhibited a slightly distinct transcriptional profile, including the enrichment of additional, less-characterized genes. These differences suggest the presence of intra-hepatocyte heterogeneity, potentially reflecting functional specialization or distinct metabolic states, while maintaining a coherent hepatocyte identity across both clusters. A more detailed dissection of the hepatocytes subpopulations is performed in subsequent analyses

###### Endothelial populations (Cluster 2; n = 8,514 ; 7.3%)

Cluster 2 was annotated as endothelial cells based on the strong enrichment of genes associated with vascular structure, endothelial identity, and transendothelial transport. Among the most discriminant markers, **STAB1** exhibited one of the highest classification performances (AUC > 0.92), with a high proportion of expressing nuclei within the cluster (86% vs 4.5% in other clusters). This gene encodes a scavenger receptor highly expressed in liver sinusoidal endothelial cells (LSECs), where it mediates endocytosis and clearance functions. Similarly, **VWF** (AUC > 0.92; 84.6% vs 1.4%) showed strong and specific expression, supporting endothelial identity through its well-established role in hemostasis and platelet adhesion. The enrichment of **STAB2** (AUC > 0.82; 64.8% vs 0.8%), another scavenger receptor specific to sinusoidal endothelial cells, further reinforces the liver-specific endothelial signature of this cluster.

Additional highly discriminant markers included **RBMS3** (AUC > 0.89; 80.6% vs 4.9%) and **FNBP1** (AUC > 0.86; 80.4% vs 17.5%), both associated with cytoskeletal organization and membrane dynamics, consistent with the high endocytic and structural plasticity of endothelial cells. The presence of **MRC1** (AUC > 0.84; 69.2% vs 0.8%), a mannose receptor involved in antigen uptake, highlights the specialized immune and scavenging functions of hepatic endothelial cells.

Genes involved in vascular signaling and endothelial maintenance were also enriched, including **KDR (VEGFR2)** (AUC ~0.78; 55.9% vs 0.5%) and **PTPRB** (AUC ~0.79; 60.0% vs 3.1%), both central regulators of angiogenic signaling and endothelial integrity. The expression of **SPNS2** (AUC ~0.81; 63.1% vs 2.7%), involved in sphingosine-1-phosphate transport, further supports roles in vascular homeostasis and immune cell trafficking.

Other markers such as **CEMIP**, **PREX2**, **LDB2**, and **IGF1R** were also enriched (pct_in ~64–66% vs low background <5%), suggesting active regulation of extracellular matrix remodeling, intracellular signaling, and endothelial responsiveness to growth factors.

Overall, the combination of highly specific endothelial and sinusoidal markers (e.g., **STAB1, STAB2, VWF, MRC1**) together with genes involved in vascular signaling and immune interactions supports the robust annotation of cluster 2 as liver endothelial cells.

###### Immune cell populations (Clusters 3, 4 and 9, n = 6,136, 5,393 and 450 ; 5.2%, 4.6% and 0.4%)

Clusters 3, 4, and 9 were collectively annotated as immune cells based on the enrichment of genes associated with hematopoietic lineage identity, immune signaling, and antigen-presenting functions.

Across these clusters, **PTPRC (CD45)** consistently emerged as a robust pan-leukocyte marker (cluster 3/4/9, AUC = 0.83/0.68/0.73; >50% vs. > 7%), thereby confirming their immune origin.

**Cluster 3** was characterized by the enrichment of genes involved in immune cell migration and activation including **DOCK2** (AUC = 0.78; 57.8% vs 3.9%) and **UTRN** (AUC = 0.78; 77% vs 54.5%) suggesting a highly dynamic cellular phenotype. Notably, transcriptional regulators associated with lymphoid lineage commitment and immune function were represented with **BCL11B** (AUC = 0.77; 54.8% vs 0.7%) and **RUNX1** (AUC = 0.68; 38% vs 2.4%) supporting a lymphoid, likely T cell-related identity.

Additional markers such as **ARHGAP15** and **PREX1** (AUC ~0.70–0.73; moderate pct_in (50/60%) with low background (<10%)) further support roles in cytoskeletal remodeling and immune cell motility, consistent with active immune surveillance functions.

**Cluster 4** displayed a distinct transcriptional profile enriched in genes associated with myeloid differentiation and immune regulation corresponding to Kupffer cells. Notably, **ZEB2** (AUC = 0.91; 87.4% vs 8.4%), **MAFB** (AUC = 0.81; 63.3% vs 0.8%) and **MARCO** (AUC=0.78; 56.5% vs 0.7%) are well-established regulators of monocyte/macrophage differentiation. The presence of **LYN** (AUC = 0.80; 64.6% vs 6.8%), a Src-family kinase involved in immune receptor signaling, and **TNFAIP2** (AUC = 0.79; 58.9% vs 1.2%), associated with inflammatory responses, further supports the identification of this cluster as a myeloid-like population. Additional genes such as **ITSN1** and **SIPA1L2** suggest active membrane trafficking and signaling processes.

**Cluster 9** was characterized by the strong enrichment of genes involved in antigen presentation and dendritic cell-like functions, including **IRF8** (AUC = 0.89; 80.4% vs 7.2%) and **FLT3** (AUC = 0.87; 73.6% vs 0.3%), both key regulators of dendritic cell development. The expression of **WDFY4** (AUC = 0.88; 78.4% vs 3.1%), associated with antigen processing, and **INPP5D (SHIP1)** (AUC = 0.85; 74.2% vs 5.1%), a negative regulator of immune signaling, further supports this identity. Additional markers such as **VAV3** and **HDAC9** suggest active intracellular signaling and transcriptional regulation within this population.

Despite these distinct transcriptional signatures, all three clusters shared a common immune identity and were therefore grouped into a broader immune compartment at this stage of the analysis. The observed differences in marker gene expression and functional enrichment suggest the presence of multiple immune subtypes, including lymphoid-like (cluster 3), myeloid/macrophage-like (cluster 4), and dendritic-like populations (cluster 9). However, given the resolution used here, these clusters were conservatively considered together, with a more detailed dissection of immune subpopulations performed in subsequent analyses.

###### Stellate cells (Cluster 6; n = 3,928 ; 3.4%)

**Cluster 6** was annotated as stellate cells based on the enrichment of genes associated with mesenchymal identity, contractile functions, and extracellular matrix organization. Among the most discriminant markers, **MYH11** showed very high classification performance (AUC = 0.93), with a strong proportion of expressing nuclei (86.0% vs 1.4%), consistent with its role as a smooth muscle myosin heavy chain and a key marker of contractile, perivascular cells. This contractile signature was further supported by the enrichment of **CALD1** (AUC = 0.84; 73.3% vs 5.7%), which encodes caldesmon, a regulator of actin–myosin interactions.

Additional genes involved in cytoskeletal organization and cellular structure were highly enriched, including **FLNB** (AUC=0.88, 83.7% vs 19.7%) and **DST** (AUC=0.86, 84.3% vs 23.4%), both contributing to mechanical stability and intracellular architecture. The presence of **NEBL** (AUC = 0.90; 81.3% vs 5.1%), associated with actin filament organization, further supports a structurally specialized and contractile phenotype.

Cluster 6 also exhibited enrichment of genes linked to extracellular matrix production and remodeling, such as **COL5A1** (AUC = 0.83; 67.3% vs 0.6%), supporting a role in matrix deposition and tissue organization, which are hallmark features of hepatic stellate cells. In addition, **FGFR2** (AUC = 0.87; 77.0% vs 2.8%) suggests responsiveness to growth factor signaling pathways involved in tissue repair and fibrogenesis.

Several additional markers, including **CCDC141**, **NRXN1**, and **SEMA5B** (all AUC > 0.86; pct_in > 70% with low background <2%), further highlight active roles in cell–cell interactions, signaling, and cellular remodeling.

Overall, the combination of strong contractile markers (**MYH11, CALD1**), cytoskeletal components (**FLNB, DST, NEBL**), and extracellular matrix-associated genes (**COL5A1**) supports the robust annotation of cluster 6 as stellate cells. The coordinated expression of these genes is consistent with a perivascular, mesenchymal population involved in maintaining liver architecture and contributing to matrix remodeling. The relatively homogeneous expression profile suggests a coherent stellate cell population at this resolution, although the presence of signaling and structural genes may reflect varying functional states.

###### Cycling and proliferative populations (Cluster 7; n = 2,539 ; 2.2%)

Cluster 7 was annotated as proliferating cells based on the strong enrichment of genes involved in DNA replication, chromosome segregation, and mitotic progression, rather than on a lineage-specific functional program. Integration of cell cycle scoring with UMAP visualization (Supplementary Figure 3 ) confirms the proliferative status of these nuclei.

Among the most discriminant markers, **BRCA1** showed the highest classification performance (AUC = 0.85), with expression detected in 70.4% of nuclei in cluster 7 compared with only 0.8% in the remaining clusters. This gene plays a central role in DNA damage repair and genome stability during cell proliferation. Other highly enriched markers included **DIAPH3** (AUC = 0.81; 61.9% vs 0.8%), **SMC4** (AUC = 0.81; 65.1% vs 4.5%), **TOP2A** (AUC = 0.80; 60.1% vs 0.8%), and **POLA1** (AUC = 0.79; 65.6% vs 10.2%). Together, these genes are associated with chromosome organization, DNA replication, and mitotic progression, supporting the interpretation of this cluster as a proliferative compartment.

This signature was further reinforced by the enrichment of additional genes involved in cell division, including **SMC2** (AUC = 0.75; 51.0% vs 0.3%), **CENPE** (AUC = 0.75; 50.8% vs 0.4%), **RIF1** (AUC = 0.74; 51.7% vs 5.3%), and **UHRF1** (AUC = 0.72; 46.4% vs 2.2%).

Importantly, the top markers of cluster 7 were mainly associated with cell proliferation and did not reveal a clear lineage-specific identity. This suggests that cluster 7 primarily reflects a shared transcriptional state linked to cell-cycle activity rather than a distinct liver cell type. Accordingly, this population was conservatively annotated as proliferating cells, likely comprising cycling nuclei originating from one or several hepatic compartments as suggested by the UMAP representation in Figure 1.

###### Cholangiocytes (Cluster 8; n = 1,083 ; 0.9%)

Cluster 8 was annotated as cholangiocytes based on the enrichment of genes consistent with a biliary epithelial identity and with duct-associated signaling functions. Among the most discriminant markers, **PKHD1** (AUC = 0.93, 85.9% vs 0.5%) and **FGFR2** (AUC = 0.91, 84.7% vs 4.5%) emerged as the most reliable indicators, showing both high specificity and high expression frequency. **PKHD1** gene encodes fibrocystin/polyductin, a protein classically associated with biliary and renal tubular epithelia, and its strong enrichment is consistent with a cholangiocyte identity. **FGFR2** is of clearer biological relevance, as fibroblast growth factor signaling has been implicated in biliary development, epithelial maintenance, and ductal remodeling, thereby reinforcing the assignment of this population to the biliary compartment.

In addition, **SHANK2** (AUC = 0.91, 82.2% vs 1.2%) was enriched in this cluster although it is not a conventional cholangiocyte marker and may reflect the existence of a distinct transcriptional program for biliary state. Several additional genes further supported the individuality of this cluster, including **GDPD5** (AUC = 0.86, 87.3% vs 44.3%), **DMD** (AUC = 0.86, 79.6% vs 18.9%), and **MAGI1** (AUC = 0.82, 83.6% vs 46.6%). Although these genes are not strictly cholangiocyte-specific, they are consistent with an epithelial program involving membrane organization, cell architecture, and signal transduction. Similarly, **ANXA5** (AUC = 0.81, 72.8% vs 22.8%), **KCNMA1** (AUC = 0.80, 68.1% vs 6.7%), and **SORBS2** (AUC = 0.80, 82.3% vs 48.9%) suggest active regulation of membrane dynamics, ion transport, and cytoskeletal organization, all of which are compatible with the polarized and specialized functions of biliary epithelial cells.

Although canonical cholangiocyte markers were not among the strongest discriminants in this dataset, **SOX9** was nevertheless enriched in cluster 8 (AUC = 0.62, 24.5% vs 0.3%), providing additional support for a biliary epithelial identity. At this resolution, the transcriptional profile appears relatively homogeneous, consistent with a discrete biliary epithelial population.

###### Megakariocytes (Cluster 10; n = 442 ; 0.4%)

Cluster 10 was annotated as megakaryocytes based on the enrichment of genes associated with platelet lineage identity, cytoskeletal organization, and platelet activation. In Figure 1, this cluster formed a small and clearly distinct population, consistent with the low abundance of megakaryocytes in liver tissue.

Among the most discriminant markers, **ITGA2B** showed the strongest classification performance (**AUC = 0.91**), with expression detected in 81.4% of nuclei in cluster 10 compared with only 0.8% in the remaining clusters. This gene encodes the integrin alpha-IIb chain, a canonical marker of the megakaryocyte/platelet lineage, and its strong enrichment provides robust support for the assignment of this cluster.

Additional markers further supported this identity. **TBXAS1** (AUC = 0.75; 55.0% vs 7.0%) encodes thromboxane A synthase 1, a key enzyme in platelet activation and hemostatic signaling, while **TFPI** (AUC = 0.73; 46.2% vs 0.3%) is involved in the regulation of coagulation pathways. The enrichment of these genes is consistent with a platelet-producing lineage specialized in hemostatic functions.

Cluster 10 also exhibited strong expression of genes related to cytoskeletal remodeling and adhesion, including **MYH9**(**AUC = 0.73**; **70.1% vs 31.3%**), **VCL** (AUC = 0.73; 52.3% vs 8.1%), and **LIMS1** (AUC = 0.76; 70.6% vs 27.8%). These genes are involved in actomyosin contractility, focal adhesion, and structural organization, which are central processes in megakaryocyte maturation and platelet biogenesis. In the same direction, **RASA3** (AUC = 0.73; 49.5% vs 4.5%) and **PLEK** (AUC = 0.68; 38.9% vs 2.8%) support an active signaling program related to platelet activation and membrane dynamics.

Several additional enriched genes, such as **KCNMA1** (AUC = 0.78; 64.0% vs 7.0%), **VAV3** (AUC = 0.74; 69.7% vs 31.9%), and **PRKD1** (AUC = 0.70; 46.2% vs 8.1%), suggest the presence of signaling and ion-regulatory mechanisms compatible with a mature and functionally specialized megakaryocytic state. Although some of these genes are not exclusive to megakaryocytes, their coordinated enrichment within this small, well-defined cluster reinforces its interpretation as a platelet-lineage population.

The distinct localization of this cluster in the UMAP and its coherent transcriptional profile are consistent with a rare but well-defined megakaryocytic population in the adult chicken liver.

Overall, the integrated analysis enabled the identification of the major liver cell populations, including hepatocytes, endothelial cells, immune cells, stellate cells, cholangiocytes, proliferating cells, and a minor megakaryocyte population. Each cluster was supported by a coherent and specific transcriptional signature, with high discriminatory power of marker genes (AUC-based) and consistent expression patterns across visualization methods (UMAP, violin plots, and heatmaps; Figure 1). While most clusters corresponded to well-defined cell types, some compartments—particularly hepatocytes, endothelial, and immune cells—showed indications of internal heterogeneity, as reflected by subtle transcriptional differences or the coexistence of multiple functional programs. To further resolve this heterogeneity and refine cell-type annotation, we next performed dedicated subclustering analyses focusing on these major populations.
